# Neural mechanisms of costly helping in the general population and mirror-pain synesthetes

**DOI:** 10.1101/2023.03.09.531639

**Authors:** Kalliopi Ioumpa, Selene Gallo, Christian Keysers, Valeria Gazzola

## Abstract

Helping others often comes with a cost to ourselves. It has been argued that experiencing the pain of others motivates helping. Here we investigate how individuals that report somatically feeling the pain of others (mirror-pain synesthetes) differ from those that do not, when deciding to help and reduce someone’s pain conveyed through different modalities. Mirror-pain synesthetes and participants who do not report such everyday life experiences witnessed a confederate expressing pain and could decide to reduce the intensity by donating money. Measuring brain activity using fMRI confirmed our initial hypothesis: self-reported mirror-pain synesthetes increased their donation more steeply, as the intensity of the observed pain increased, and their somatosensory brain activity (in SII and the adjacent IPL) activity was more tightly associated with donation when the pain of other was conveyed by the reactions of the pain-receiving hand. For all participants, in a condition where the pain was conveyed by facial expressions, activation in insula, SII and MCC correlated with the trial by trial donation made, while SI and MTG activation was correlated with the donation in the Hand condition. These results further inform us about the role of empathy in costly helping, the underlying neural mechanism, and individual variability.

## Introduction

To help someone in need we often need to sacrifice something ourselves. It has been proposed that feeling the pain of others as if it were our own is a key motivator to help. This idea was brought to prominence through Adam Smith’s theory of moral sentiments (1759): “As we have no immediate experience of what other men feel, we can form no idea of the manner in which they are affected, but by conceiving what we ourselves should feel in the like situation. Though our brother is upon the rack, […] it is by the imagination only that we can form any conception of what are his sensations. Neither can that faculty help us to this any other way, than by representing to us what would be our own, if we were in his case. It is the impressions of our own senses only, not those of his, which our imaginations copy. By the imagination we place ourselves in his situation, we conceive ourselves enduring all the same torments, we enter as it were into his body, and become in some measure the same person with him, and thence form some idea of his sensations, and even feel something which, though weaker in degree, is not altogether unlike them. His agonies, when they are thus brought home to ourselves, when we have thus adopted and made them our own, begin at last to affect us, and we then tremble and shudder at the thought of what he feels. For as to be in pain or distress of any kind excites the most excessive sorrow, so to conceive or to imagine that we are in it, excites some degree of the same emotion, in proportion to the vivacity or dullness of the conception”. The notion that empathy promotes prosociality has received empirical support (Batson et al., 1981; FeldmanHall et al., 2015; Jordan et al., 2016; Smith, 1759 but see Vachon et al., 2014), but what is exactly meant by “enduring all the same torments” however remains somewhat unspecified: the subjective experiences of witnessing the pain of others differs across individuals, with some merely experiencing emotional distress while others experience localized somatic feelings broadly matching those observed. Specifically, some report feeling tactile sensations on their own skin when observing touch on others (mirror-touch synaesthesia) or report somatic pain in their own body while observing the pain of others (mirror-pain synesthesia/vicarious pain perception) (Banissy et al., 2009; Banissy and Ward, 2007; Blakemore et al., 2005; Fitzgibbon et al., 2010). Whether such added somatic feelings influence the motivation to help is at the center of the present study, and tests Smith’s intuition that it is our own sensory experiences that are key to excite our own emotion and sympathy.

Behaviorally, existing studies link mirror-sensory synesthesia with (i) enhanced empathy as assessed by questionnaires (emotional reactivity in Banissy and Ward, 2007 and Ward et al., 2018, empathic concern in Ioumpa et al., 2019) (ii) enhanced empathic accuracy while recognising subtle facial expressions (Banissy et al., 2011; Ward et al., 2018) and (iii) increased self-report affect intensity when looking at emotional pictures (Ioumpa et al., 2019). These effects seem restricted to the affective dimension of empathy, as synesthetes do not seem to have enhanced ‘theory of mind’ cognitive empathy skills as measured by the Reading the Mind in the Eyes test (Baron-Cohen et al., 2016) and the ‘movie for the assessment of social cognition’ (MASC) test (Santiesteban et al., 2015). Also synesthetes seem to score lower on the social skills scale of the EQ (Baron-Cohen et al. 2016; Ward et al., 2018). Whether their increased affective empathy translates into increased prosociality however remains poorly understood. Encouraging evidence stems from Ioumpa et al., (2019) who found that mirror-sensory synesthetes donate more money to a stranger in a dictator game, but our core question of whether the added somatic sharing of pain in synesthesia would increase helping when witnessing the pain of others remains unexplored.

To test the impact of mirror-pain synesthesia on helping behavior, we here adapt a costly helping paradigm introduced by Gallo et al., (2018), in which participants are given the opportunity to donate money to reduce the pain of a victim they witness receive a noxious stimulation via what they believe to be a close circuit camera (Figure 1). Importantly, the pain level is conveyed either by a facial expression of pain triggered by an electric shock (Face condition) or by the kinematics of a hand being slapped by a belt (Hand condition). These two stimulus types were developed to compare conditions that should merely trigger vicarious distress (faces) from those that could encourage the somatosensory mapping onto a specific region of the observer’s body (hand) (see Gallo et al., 2018; Keysers et al., 2010). Accordingly, if somatic mapping on the observer’s body is increased in mirror-pain synesthetes, and this motivates helping, we expect mirror-pain synesthetes to increase their donations more steeply when observing more pain particularly in the hand stimuli.

**Figure 1.**
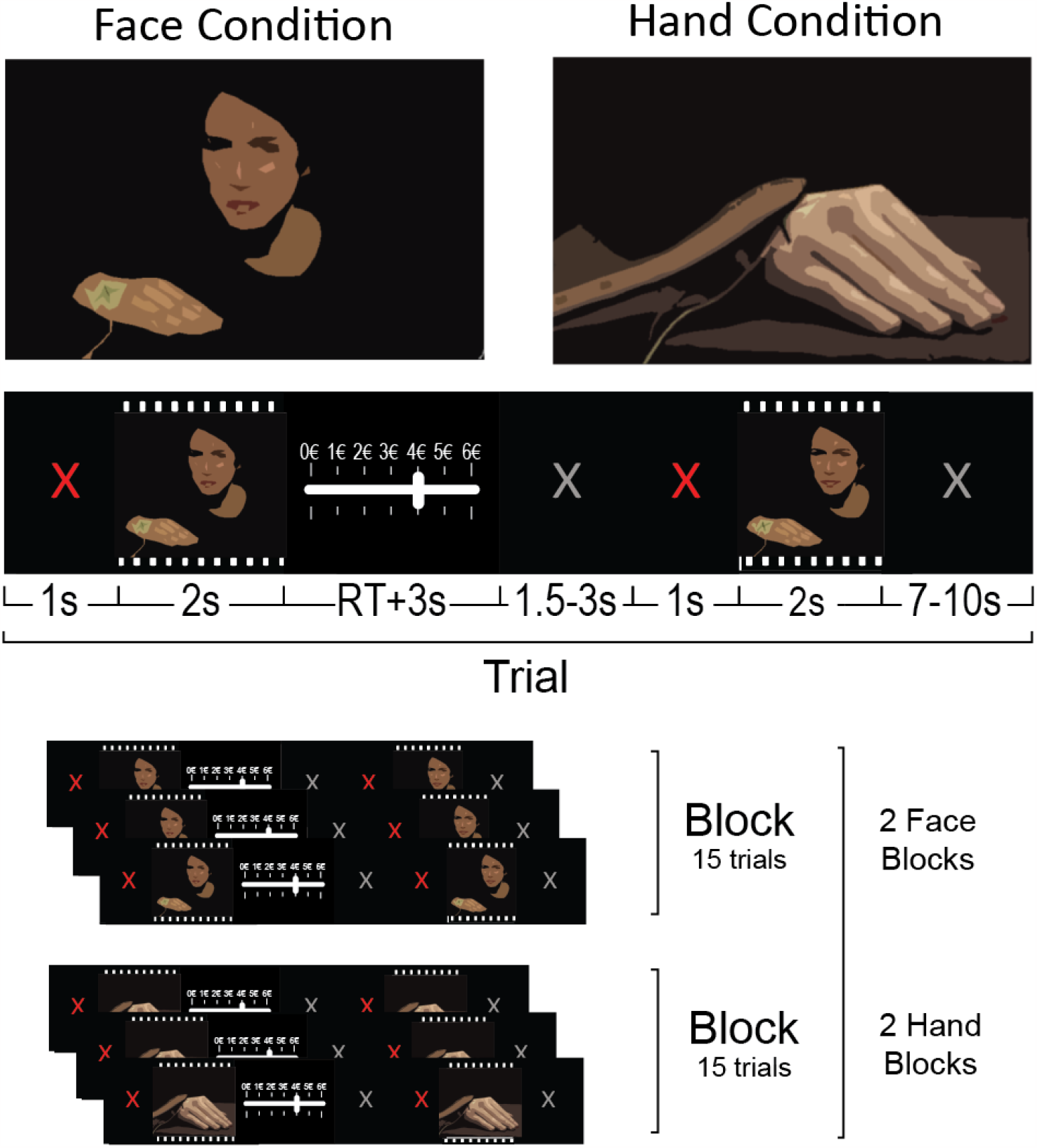
Helping task structure. Top: two screen shots taken from a video showing the confederate receiving an electrical shock on the hand and manifesting its painfulness through facial reactions and a video showing the confederate receiving a slap on her right hand. Middle: trial structure. Bottom: the experiment consisted of two Face and two Hand blocks were presented of 15 trials each.

Neurally, first-person experiences of pain are thought to result from the combination of a sensory-discriminative dimension (where and what kind of pain do I feel?) and an affective dimension (how aversive is this pain?), with the former associated with activity in somatosensory cortices (SI and SII) while the latter is associated with activity in the anterior insula and cingulate cortex (Keysers et al., 2010; Mouraux et al., 2011; Price, 2000). In agreement with the general notion that empathy reflects mapping the pain of others onto our own pain, brain regions and neurons involved in our own pain are activated while witnessing the pain of others (Carrillo et al., 2019; de Waal and Preston, 2017; Keysers et al., 2010; Lamm et al., 2011; Singer et al., 2004). That this mirroring of the affective component of pain may promote prosociality is borne out by evidence that activations in the affective pain areas correlates with helping someone in pain (Christov-Moore and Iacoboni, 2016; FeldmanHall et al., 2015; Hein et al., 2010; Ma et al., 2011; Tomova et al., 2016) and inhibiting regions involved in pain experience, the cingulate in particular, reduces helping in rodents (Hernandez-Lallement et al., 2020).

Distinguishing affective from sensory components refines this picture: witnessing the pain of others independently of how it is perceived activates the more affective brain regions (rostral cingulate and anterior insula in particular), while witnessing the details of how a specific body part is harmed additionally triggers activity in SI or SII (Ashar et al., 2017; Bufalari et al., 2007; Christov-Moore and Iacoboni, 2016; Decety, 2011; Keysers et al., 2010; Keysers and Gazzola, 2009; Krishnan et al., 2016; Lamm et al., 2011; Morrison et al., 2013; Nummenmaa et al., 2012; Shih et al., 2008; Singer and Lamm, 2009), and altering activity in SI alters helping (Gallo et al., 2018). Mapping the somatic feelings reported by mirror-pain synesthetes onto this distinction would suggest that they may activate SI and SII more while witnessing the sensations and pain of others. Indeed, mirror-pain synesthetes have higher activity (Osborn and Derbyshire, 2010) and increased gray matter (Grice-Jackson et al., 2017) in somatosensory cortices and anterior insula compared to controls participants; and Blakemore et al. (2005) found enhanced SI and SII activation when one mirror-touch synesthete watched movies of other people being touched compared to control participants. Hence, we might expect participants that report experiencing mirror-pain synesthesia to show more SI/SII activation when witnessing a hand being slapped, and if this somatic sharing indeed contributes to the motivation to help, this SI or SII activity should be more tightly associated with the amount of money donate to reduce that pain in mirror-pain synesthetes than non-synesthetes when witnessing the belt hitting a hand.

Additionally, recent studies have identified multivariate brain patterns that are somewhat selectively recruited when participants experience (i) physical pain (wager et al., 2013), (ii) the feeling of guilt (Yu et al., 2020), and (iii) witness other people’s pain (Krishnan et al., 2016; Zhou et al., 2020). We might expect that while all these patterns could be associated with the motivation to help (and hence the amount of money donated in our task), a pattern trained to decode physical pain should be most tightly associated with donation in participants reporting mirror-pain synesthesia.

To shed light on the contribution of vicarious somatic pain as a motivator of helping, and test the above-mentioned hypotheses, here we therefore recruited participants that report experiencing mirror-pain synesthesia and some that do not, and measured their willingness for costly helping other individuals (as in Gallo et al., 2018) while also measuring their brain activity using fMRI.

## Materials and Methods

### Participants

In total, 32 healthy volunteers (37y±17SD; 32f) with normal or corrected-to-normal vision, and no history of psychiatric, neurological, other medical problems, or any contraindication to fMRI participated in our experiment. Participants were recruited through advertisements of the experiment on social media advertising a helping decision-making study (25 participants). In addition, we also invited individuals with mirror-pain synesthesia through the contact list of participants from the study by Ioumpa et al., 2019 where they had taken part as synesthetes (7 participants).

In the end of the experiment all participants were asked to provide a “Yes-No” answer to whether they have mirror-pain synesthesia experiences during their everyday life (*“In mirror-pain synesthesia people feel on their own body the pain they observe in others. Do you have such experiences in your everyday life?”*). Those who reported having everyday mirror-pain synesthesia-like experiences were classified as self-report mirror-pain synesthetes (13 participants) and are the focus of the paper. Participants who did not report experiencing mirror-pain synesthesia experiences in their everyday life will be referred to as control participants in this study. As an additional quality check measure, all participants filled the Vicarious Pain Questionnaire developed by Grice-Jackson et al., (2017). This classification revealed 7 sensory/localiser and 3 affective/general participants in our sample while the rest of our participants were classified as non-responders. Importantly, being classified as sensory/localiser was 8 times more likely amongst the self-reported mirror touch synesthetes than amongst those not self-reporting mirror touch synesthesia (Table S1). Due to the small group sizes resulting from this finer classification, we created a responders (sensory/localiser and affective/general participants together) and no responders group, and only used this tool for our analyses. A more detailed description can be found in Supplementary information S1) As two (who were recruited as synesthetes) of the 32 participants were left handed, and stimuli showed movements of the right hand of the actor, in order to reduce potential variability induced by lateralization of the brain responses, these two participants only performed the tasks off-line (i.e. no fMRI data acquired). One control fMRI participant was excluded for having a very low correlation (<0.2) between video intensity (as given from an initial stimuli validation) and donation, for both the Hand or Face conditions. This was a criterion that we had set from the beginning for inclusion in our Helping Paradigm analyses. Thus we ended up with fMRI data for 29 participants (11 self report mirror-pain synesthetes and 18 control participants) and behavioral data for 31 participants (13 self report mirror-pain synesthetes and 18 control participants). The study was approved by the Ethics Committee of the University of Amsterdam, The Netherlands (2017-EXT-8201). Consent forms for participation and authorization for the publication of images have been obtained.

### fMRI Helping Paradigm

#### Stimuli

The same set of stimuli as in Gallo et al. 2018 was used. Two types of videos were presented. One showing the confederate receiving an electroshock on the hand and expressing the pain she felt by only reacting with facial expression (Face videos). The other showed a belt hitting the dorsum of the confederate right hand, and the confederate expressing how much pain she felt by a reaction of the hand alone (Hand videos). The face was not visible in the latter stimuli. All videos lasted 2s and were neutral during the first second. The Face videos started with the face in a neutral expression that was kept neutral until the stimulation. The Hand videos started with the belt laying on the hand dorsum until the end of the first second when it was lifted in order to hit. Stimuli were validated by an independent group of subjects (45 volunteers, 32.36y±10.36SD; 23f), who were asked to rate the intensity of the pain displayed on a scale from 1 to 10, with ‘1’ being ‘just a simple touch sensation’ and ‘10’ being ‘most intense imaginable pain’. Videos were edited using Adobe Premiere Pro CS6 (Adobe, San Jose, CA, USA).

#### Task

The task was an fMRI adaptation of the Helping task as published in Gallo et al. (2018). Participants performed 60 trials in which they watched a first (pre-recorded) video of the confederate receiving a painful stimulation. The intensity of the stimulation could vary between 1 and 6 on a 10 point pain scale, and was chosen on each trial randomly by the computer program. In each trial participants received 6 euro credits, and could decide to donate some of them in order to reduce the intensity of the second stimulation to the confederate. Each donated credit reduced the next stimulation by 1 point on the 10 point pain scale. Participants then watched a second video showing the confederate’s response to the second stimulation. At the end of the task, participants were paid the sum of the amount of money that they had kept for themselves from all the trials divided by 10. Prosocial behavior was captured as the average number of credits given up in all trials (“donation”). Hand and Face videos were presented in separate sessions that were randomized across participants. In total there were 2 sessions of 15 trials for the face and hand videos.

We used the same cover story used in Gallo et al. 2018. Each participant was paired with what they believed to be another participant like them, although in reality it was a confederate. They drew lots to decide who plays the role of the decision maker and of the pain-receiver. The lots were rigged so that the confederate would always be the pain-receiver. The participant was then taken to the scanning room while the confederate was brought to an adjacent room, with a fake filming set up. Participants were misled to believe that the pain stimulations were delivered to the confederate and displayed to them in fMRI in real-time while in reality pre-recorded videos were used. All participants were presented the same set of videos in a randomized order.

At the end of the fMRI tasks, participants were debriefed. To assess whether they believed the cover story, they were asked to answer the question ‘Do you think the experimental setup was realistic enough to believe it’ on a scale from 1 (strongly disagree) to 7 (strongly agree) in an exit questionnaire. All participants reported that they at least somewhat agreed with the statement (i.e. 5 or higher). Participants were also asked to fill out the interpersonal reactivity index (IRI) empathy questionnaire (Davis, 1983), and the money attitude scale (Yamauchi and Templer, 1982).

The task was programmed in Presentation (www.neurobs.com), and presented under Windows 10 on a 32 inch BOLD screen from Cambridge Research Systems visible to participants through a mirror (distance eye to mirror: ∼10cm; from mirror to the screen: ∼148cm). The timing of the task was adapted to the requirements of fMRI: Each trial started with a jittered gray fixation cross lasting 7-10 seconds (Figure 1). Then a red fixation cross appeared for 1 second, followed by the first video presentation and the donation scale. Participants could make their choice without a time restriction. In order to make their choice they could move the bar in the scale using their right index and middle finger. After 3 seconds of inactivity the system would automatically register their response. Then a jittered gray fixation cross lasting 1.5-3 seconds would follow, then a 1 second red cross and the second video. The role of the red fixation crosses was to capture participants’ attention just before a video appears.

### Analysis of Behavioral Data

Statistical analyses were performed using JASP (https://jasp-stats.org, version 0.11.1), to provide both Bayes factors and *p* values. Bayes factors allow us to differentiate between evidence of absence and evidence of the presence of an effect, and therefore complement traditional frequentist statistics as p-values cannot quantify evidence for the absence of an effect (Keysers et al., 2020). We used traditional bounds of BF_10_>3 to infer the presence of an effect and BF_10_<⅓ to infer the absence of an effect (Keysers et al., 2020). Two-tailed tests are indicated by BF_10_,i.e. *p*(Data|H_1_)/*p*(Data|H_0_) while one-tailed tests are indicated by BF_+0_. Where ANOVAs were used, we report BF_incl_ which reports the probability of the data given a model including the factor divided by the average probability of the data given the models not including that factor. Normality was tested using Shapiro-Wilk’s. We always used default priors for Bayesian statistics as used in JASP.

### MRI Data acquisition

MRI images were acquired with a 3-Tesla Philips Ingenia CX system using a 32-channel head coil. One T1-weighted structural image (matrix = 240×222; 170 slices; voxel size = 1×1×1mm) was collected per participant together with an average of 775.83 EPI volumes ± 23.11 SD (matrix M x P: 80 × 78; 32 transversal slices acquired in ascending order; TR = 1.7 seconds; TE = 27.6ms; flip angle: 72.90°; voxel size = 3×3×3mm, including a .349mm slice gap).

### fMRI Data preprocessing

The MRI data were processed in SPM12. EPI images were slice-time corrected to the middle slice and realigned to the mean EPI. High quality T1 images were coregistered to the mean EPI image and segmented. The normalization parameters computed during the segmentation were used to normalize the gray matter segment (1mmx1mmx1mm) and the EPIs (2mmx2mmx2mm) to the MNI templates. In the end, EPIs images were smoothed with a 6mm kernel.

### fMRI Data analyses

Our analyses and the experimental design focused on how brain activity during the first video influenced donation. The second videos were modeled, but as a variable of no interest as they were the result of a decision rather than the cause of that decision.

#### GLM analyses

For each of our four sessions (two that presented the Face stimuli and two the Hand), our fMRI design matrix included the following regressors. (1) A first video regressor that started with a red fixation cross and ended with the end of the first video. We call this regressor the main effect of Face or Hand Video1. The red cross was included in this regressor as it was always presented at a fixed time interval of 1s before the first video and separating their contribution to the BOLD signal would not have been possible. (2) This regressor had the donation made in each trial as a parametric modulator, creating a FaceDonation and a HandDonation parametric modulator. The donation values for each run were standardized with the zscore function of MATLAB before being inserted as a regressor. Standardization here was used so that the parameter estimate of the parametric modulator becomes independent of a participants range of donation, and reflects how tightly brain activity is associated with donation (in the sense of a correlation) rather than the specific slope. (3) A decision regressor started with the appearance of the donation scale and ended 3 seconds after the last button press of the participant, when the scale disappeared. (4) The second video regressor was aligned with the presentation of the red cross before the second video and ended with the end of the second video. (5) A regressor with the standardized donation made in this trial as a parametric modulator for video 2. (6-11) Finally, 6 regressors of no interest were included to model head translations and rotations.

We then brought the parameter estimate images for FaceDonation, HandDonation and Face and Hand main effects into four separate t-tests and contrasted them against zero. Results were thresholded at p_unc_ < .001 and 5% family-wised error (FWE) corrected at the cluster level by setting the minimum cluster-size k to the FEWc value calculated by SPM after visualizing the results at p_unc_<0.001 k=10 (Eklund et al., 2016).

#### Neurological signatures analyses

Because of the difficulties to associate changes in brain activity in a single location with specific mental processes without facing reverse inference issues (Poldrack, 2006), we additionally used three multivariate signatures. These maps quite selectively detect whether participants perceiving other people’s pain (vicarious pain signature, VPS, Krishnan et al., 2016), feel their own pain (neurological pain signature, NPS, Wager et al., 2013) or feel interpersonal guilt (Yu et al., 2020). In order to explore if signals in these networks covaried with FaceDonation and HandDonation we brought the signatures into our fMRI analysis space using ImageCalc, extracted the FaceDonation and HandDonation parameter estimate image (β_FaceDonation_ and β_HandDonation_) from each participant and dot-multiplied them separately with the three signatures. The result indicated how much the covariance with FaceDonation and HandDonation loads on the VPS, NPS and guilt signatures. We then brought these values into JASP, and compared them against zero and checked for differences between individuals reporting mirror-pain synesthesia experiences and those who did not.

## Results

### Behavioral results

Participants donated the same amount on average for the face (mean±SD, 2.538±1.066) and hand (mean±SD, 2.509±1.149) conditions (t_(30)_=0.289, *p*=0.774, Cohen’s d=0.052, BF_10_=0.199). Also participants donated more money on trials in which the confederate expressed more pain both for the Face (average correlation across participants r=0.816, SD=0.168) and the Hand (average correlation across participants r=0.634, SD=0.220) conditions.

Figure 2 shows that when comparing the donation made between self-report mirror-pain synesthetes and control participants we observed that synesthetes donated more money in order to help than control participants for for the Face (t_(29)_=4.719, *p*<.001, Cohen’s d=1.718, BF_+0_=692.648) and Hand conditions (t_(29)_=3.917, *p*<.001, Cohen’s d=1.426, BF_+0_=108.411).

**Figure 2.**
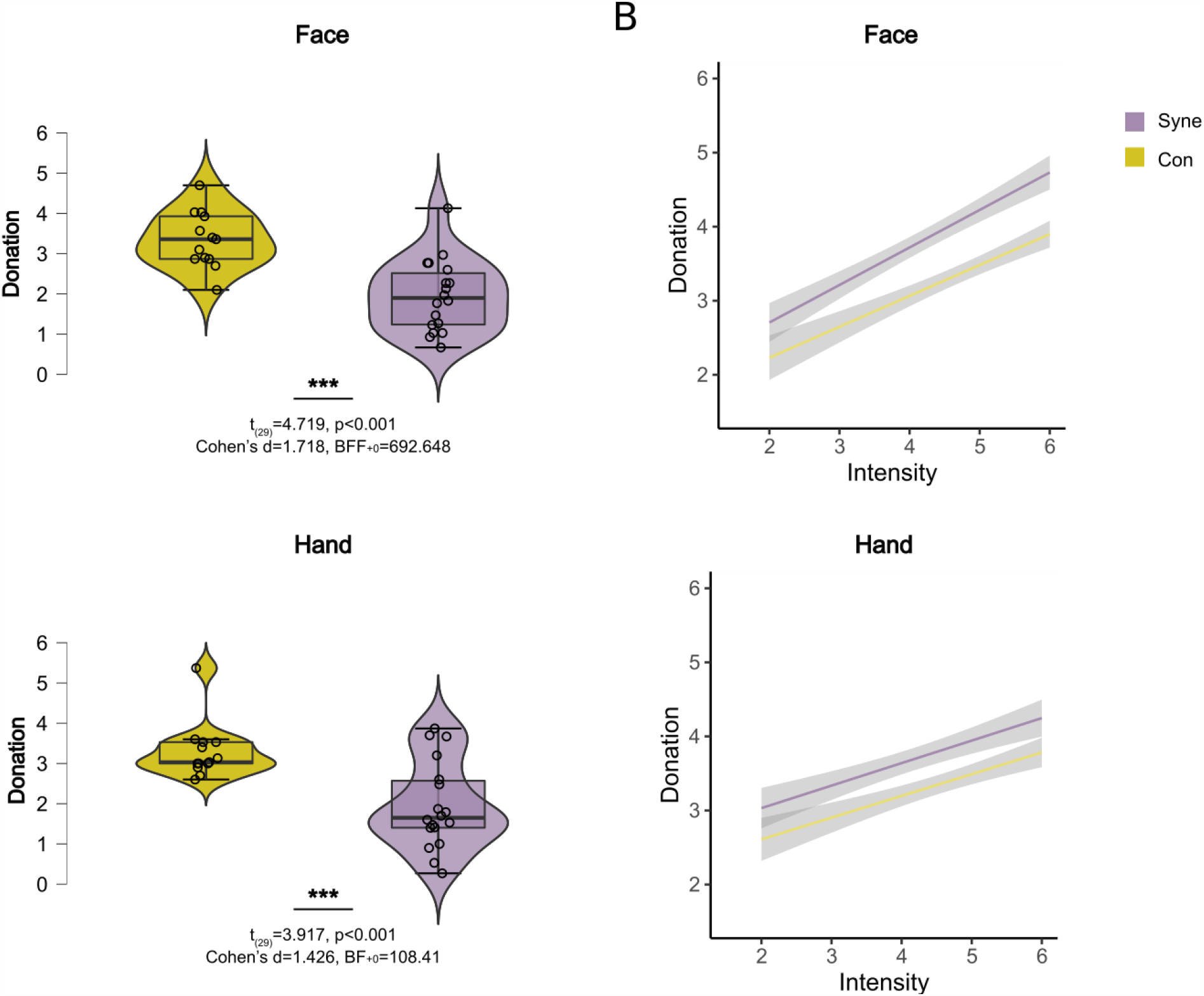
**(A)** Average of donation for the Face and Hand conditions for self-report synesthetes (cyan) and controls (yellow). Violin plots represent the distribution, the box-plot within, the median, and the whisker the quartiles. The BF_10_ and p-values between the violin plots represent the results of the comparison between individuals reporting mirror-pain synesthesia experiences and those who did not. **(B)** Correlation between Donation and Intensity for the Face and Hand conditions for self-report synesthetes and controls.

We analyzed the relationship between the intensity of video1 (as resulted from the stimuli validation described in Gallo et al. 2018) and the donations as a function of the stimulus (Hand vs Face) and self-report synesthesia using a random intercept linear mixed model with subject as random effect. Decomposing the effect of these factors revealed that participants gave 2.54 euros per trial on average, and that the donation depended most strongly on the intensity of video1 (F_(1,1764)_=1369.176, p<0.001), with a slope of 0.65. Importantly, self-declared synesthetes gave more on average (3.32 ±0.66 SD) than controls (1.94 ±0.94 SD) and also had a steeper slope (F_(1,1764)_=23.354, p<0.001), i.e. adapted their donations more to the intensity of the victims pain. This group difference did not depend on whether Face or Hand stimuli were seen (F_(1,1764)_=0.113, p<0.736).

Comparison responders and no responders groups from the VPQ we observed the same tendency with responders donating significantly more for the face (t_(26)_=-1.722, *p*=0.048, Cohen’s d=-0.679, BF_+0_=1.951) and hand (t_(26)_=-1.877, *p*=0.036, Cohen’s d=-0.740, BF_+0_=2.406) conditions when using frequentist statistics, while the Bayesian statistics were less conclusive, although showing a similar trend.

None of the subscales of IRI or MAS correlated with the donation and the Bayesian statistics were close to evidence for absence of a correlation (results at **Table S2**).

### fMRI results

#### GLM analyses

When looking at the main effect of video1, i.e. voxels where the BOLD signal is increased while viewing the first video, independently of donation, and irrespectively of whether the pain was conveyed by the facial expression or the hand movement, we observed a network resembling the pain observation network often reported in the literature, including the ACC, MCC, SII and Insula (**Supplementary Fig. S1 and Supplementary Table S3**), suggesting that witnessing a painful stimulation delivered to the confederate triggered expected neural response. Comparing the main effect of Face and Hand during the the first video revealed significant differences across these two types of stimuli: the IFG and IPL showed higher BOLD signal for Face than Hand stimuli and SII, insula and the calcarine gyrus showed higher BOLD signal for the Hand than Face (**Supplementary Fig. S2 and Supplementary Table S4**). Comparing self-declared synesthetes and controls for the main effect of Face or Hand (i.e. independently of donation) did not yield significant differences.

We then localized voxels in which activity correlated with Donation in all participants for Hand and Face trials separately. For the Face condition, we observed that the more money participants donated the higher the BOLD signal in the insula, SII, TPJ and MCC (**Fig. 3** and **Table 1**).

**Table 1.**
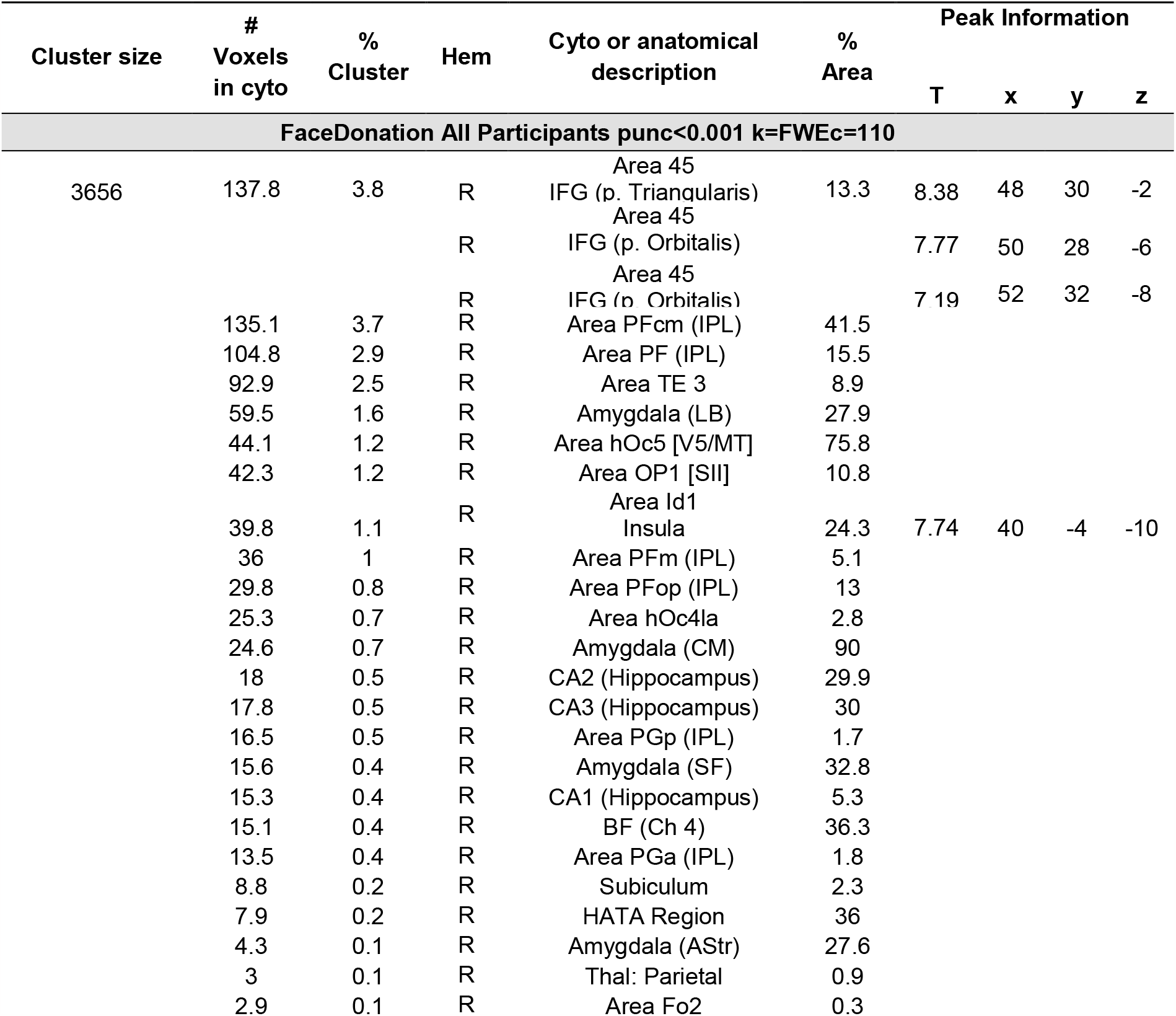
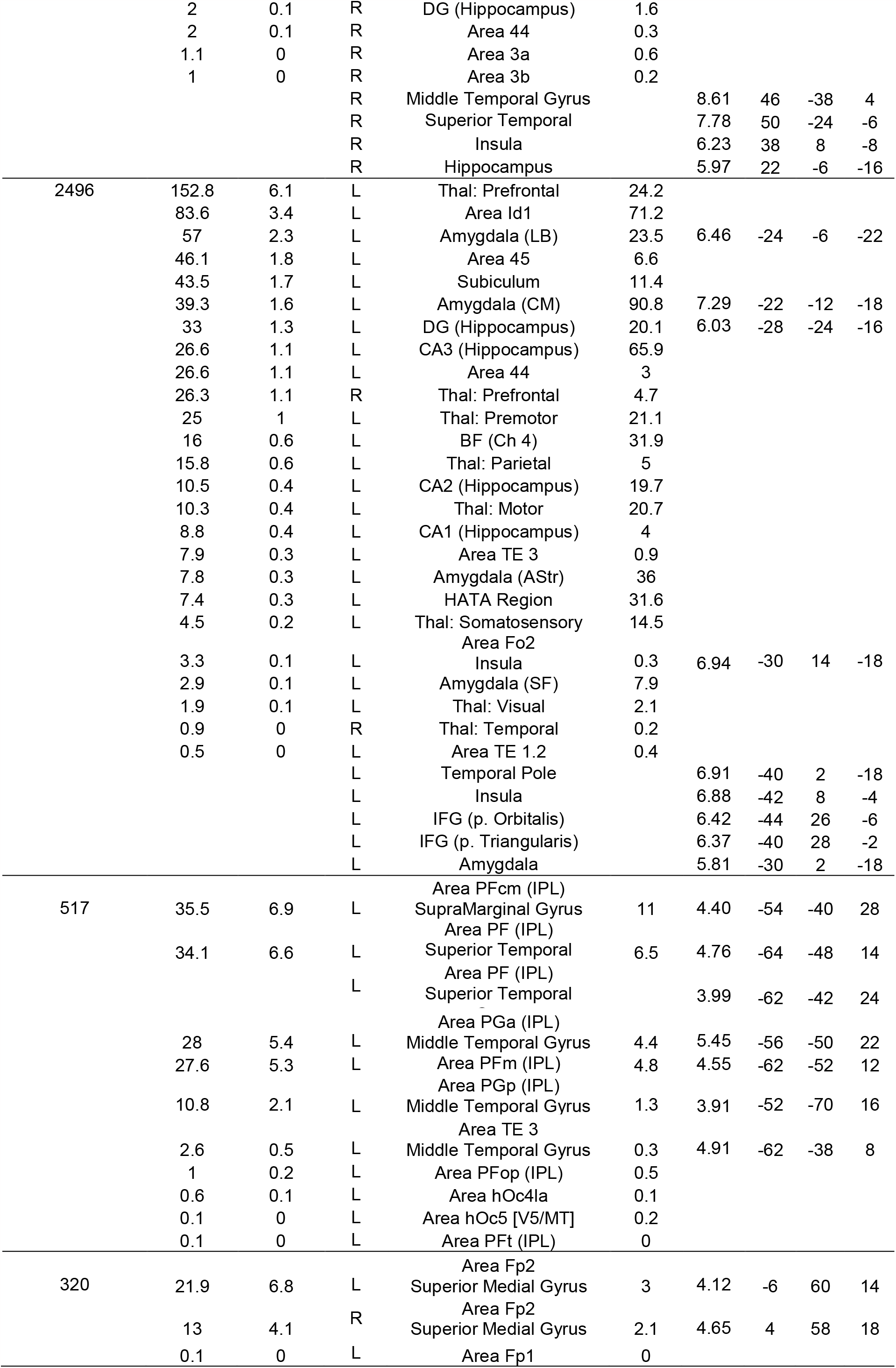
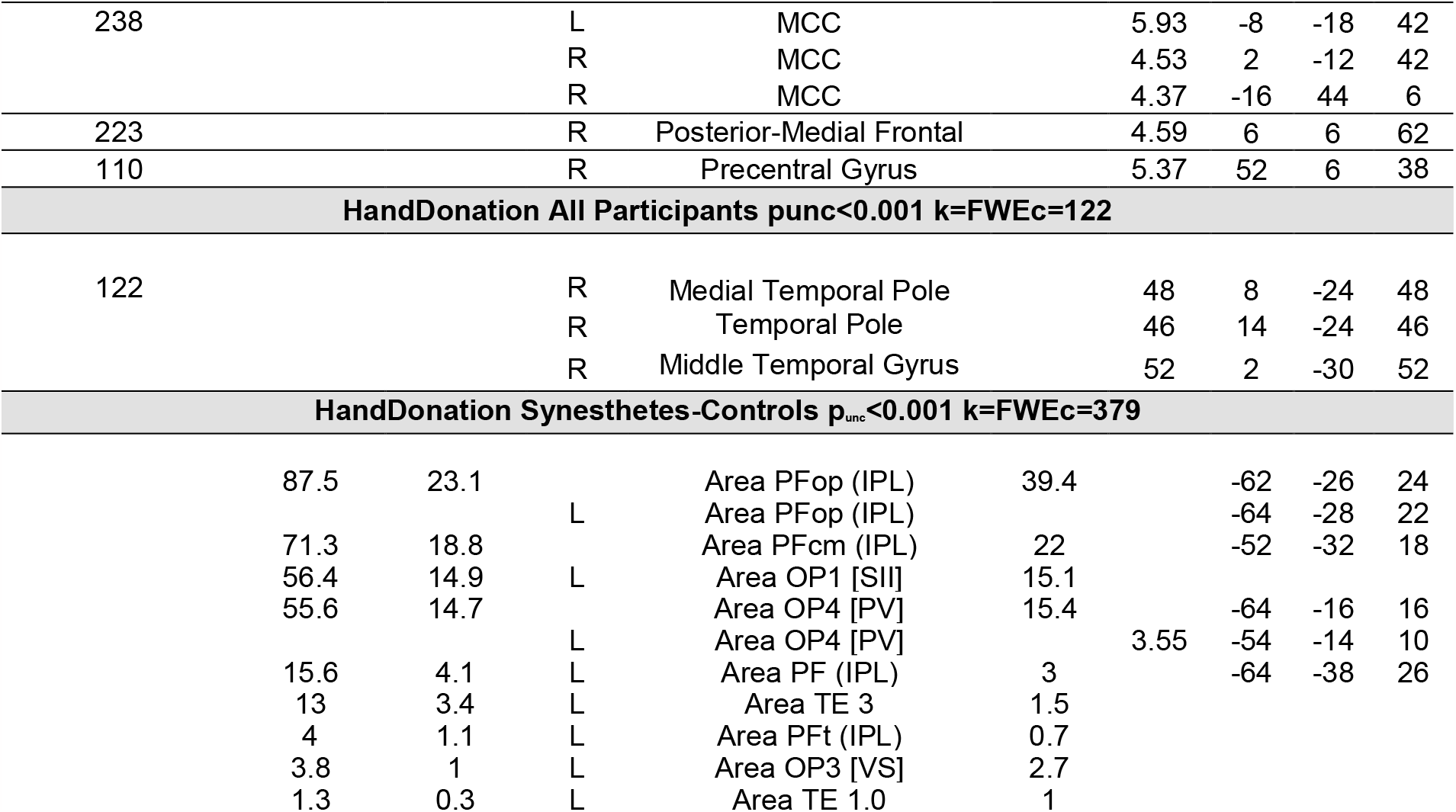
Results of the voxelwise analysis. Brain activations for the FaceDonation for all participants together, for the HandDonation for all participants and for the HandDonation for Synesthetes-Control participants. Regions were labeled using SPM Anatomy Toolbox. From left to right: the cluster size in number of voxels, the number of voxels falling in a cyto-architectonic area, the percentage of the cluster that falls in the cyto-architectonic area, the hemisphere (L=left; R=right), the name of the cyto-architectonic area when available or the anatomical description, the percentage of the area that is activated by the cluster, the t values of the peaks associated with the cluster followed by their MNI coordinates in mm.

**Figure 3.**
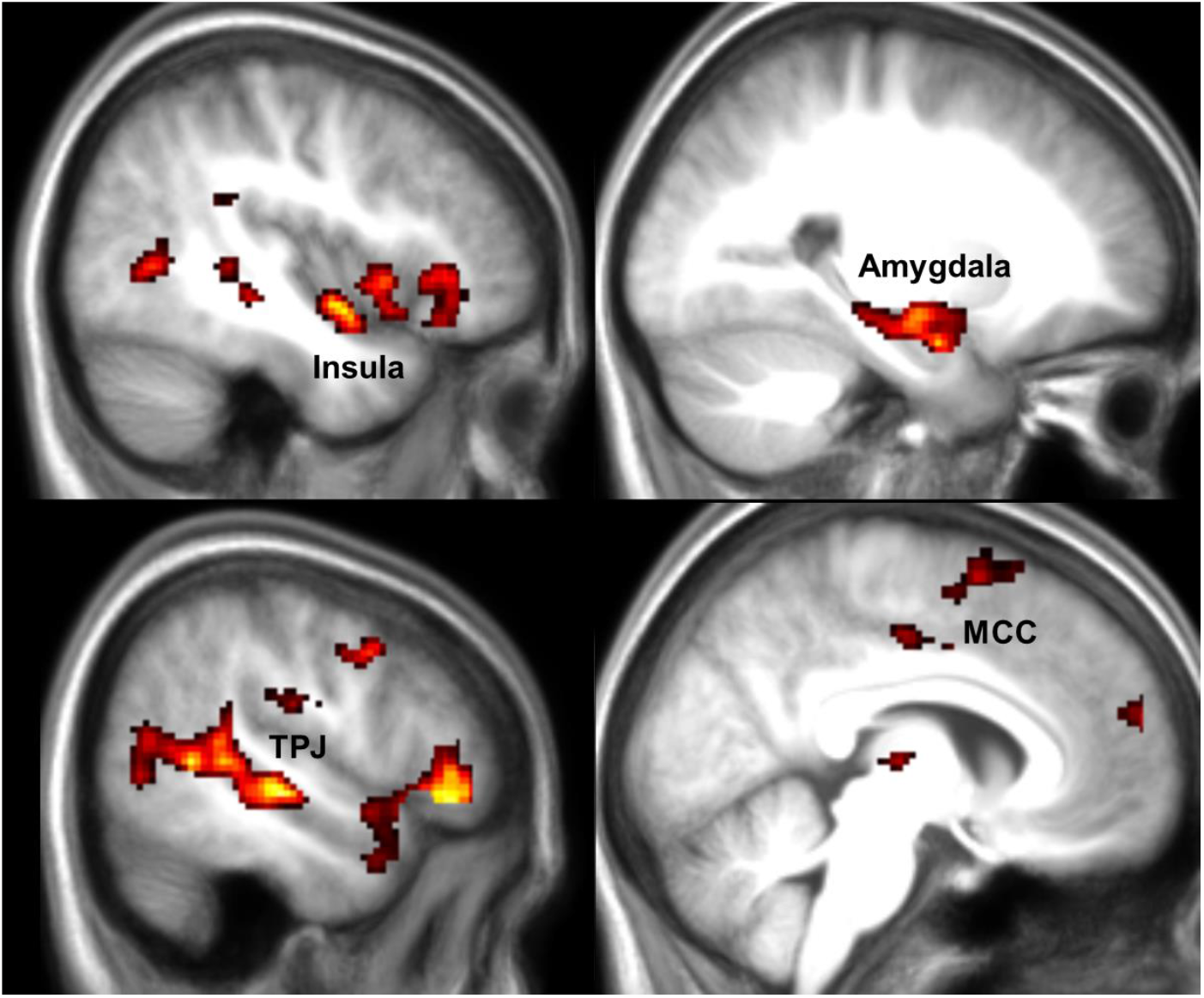
fMRI results for Face Donation. Results of a linear regression on the parametric modulator for the first video and trial-by-trial donation in the Face condition. Results are FWE cluster-corrected at p<0.05 (p_unc_<0.001, k=FWEc=110 voxels, 3.34<t<8). This identifies voxels with signals that increase for higher donation.

For Hand trials, we observed that the more money participants donated the higher the BOLD signal in the Middle Temporal Gyrus (**Fig. 4** and **Table 1**). Based on EEG results in Gallo et al. (2018) we expected SI to also predict donation in the Hand condition. We then looked at the results at uncorrected p_unc_=0.01 and found activation in SII, insula, ACC, MMC among others (**Supplementary Figure S3 and Supplementary Table S5**). Driven by the surprising lack of findings for SI even at a lower threshold we decided to use a multivariate approach (partial least-square regression) which sometimes has higher sensitivity than univariate analyses for specific regions of interest, to further explore the role of SI in the Hand condition (procedure described at **Supplementary information S3**). This multivariate approach revealed that SI does indeed contain information that relates to the magnitude of donations when a pain is conveyed by the hand (**Supplementary Figure S4**).

**Figure 4.**
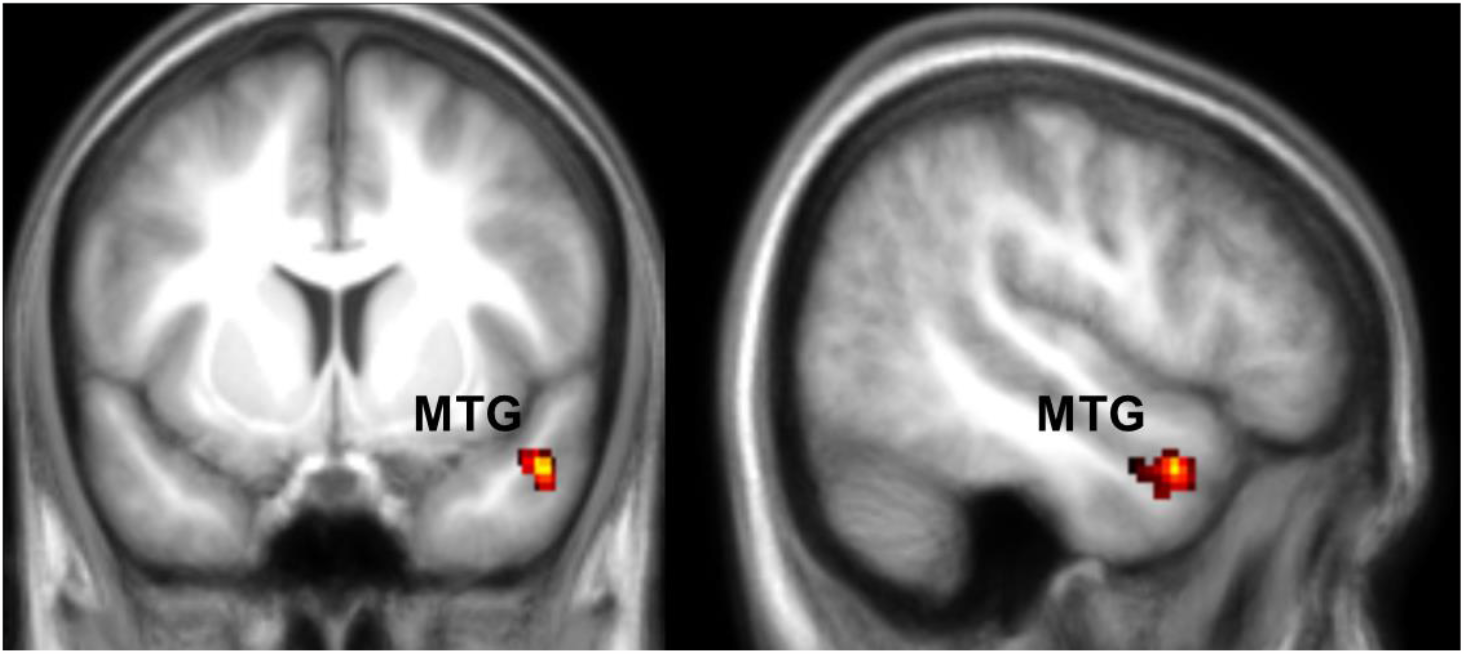
fMRI results for Hand Donation. Results of a linear regression on the parametric modulator for the first video and trial-by-trial donation in the Hand condition. Results are FWE cluster-corrected at p<0.05 (p_unc_<0.001, k=FWEc=122 voxels, 3.34<t<5). This identifies voxels with signals that increase for higher donation.

To explore the difference between the association with donation for Face and Hand trials in more detail, we directly compared the parameter estimates for the parametric donation modulators for Hand and Face using a paired sample t-test, but no significant results survived for either direction. This could suggest that a similar network is activated during the Hand condition as well but less strongly or in a more variable way across participants.

To explore whether self-report mirror-pain synesthetes differed in their brain activations compared to self-report non-synesthetes, we performed a two sample t-test comparing the FaceDonation and HandDonation parametric modulators across these two groups. We observed higher parameter estimates in SII and the adjacent parietal operculum for the self-report synesthetes compared to the non-synesthetes for the HandDonation regressor contrast (**Fig. 5** and **Table 1**). This suggests that, as hypothesized, donations are more tightly associated with somatosensory activity in synesthetes than controls. The FaceDonation comparison did not reveal significant differences.

**Figure 5.**
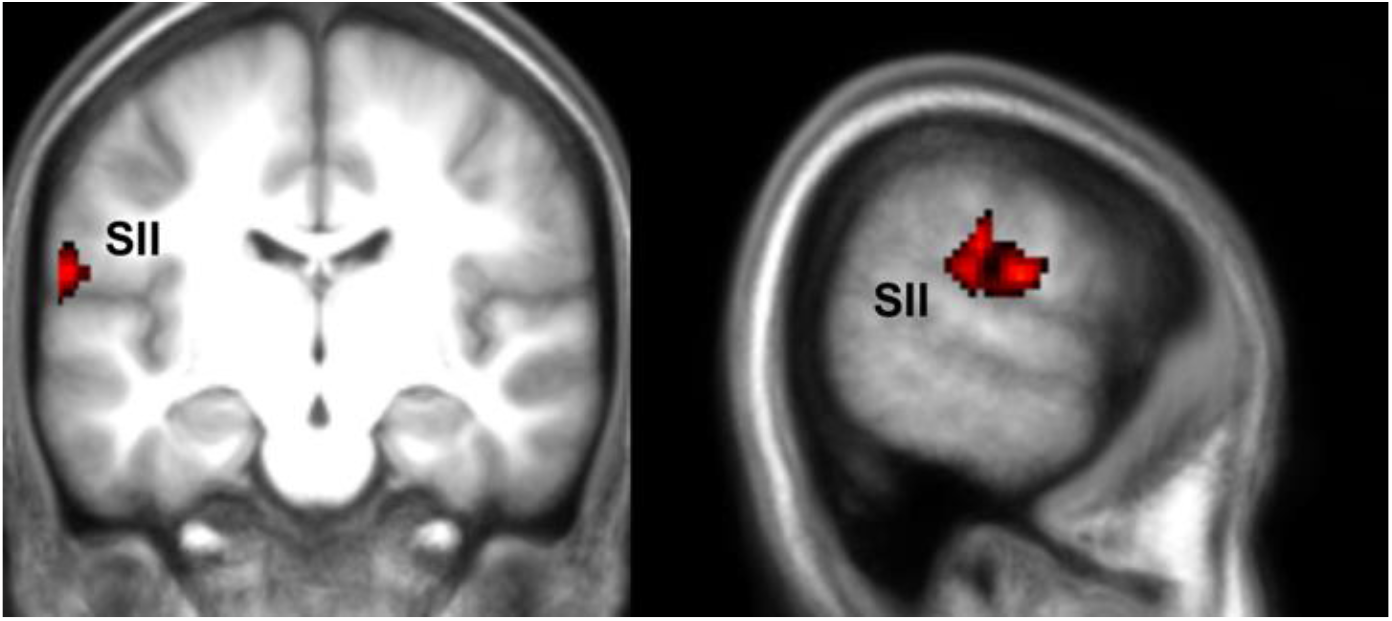
fMRI results for Hand Donation group comparison. Results of a two sample t-test between the self-report synesthetes and control participants for the HandDonation. Results are FWE cluster-corrected at p<0.05 (p_unc_<0.001, k=FWEc=379, 3.34<t<8). This identifies voxels with signals that increase for higher donation. The reverse contrasts did not yield results.

#### Neurological signatures analyses

When looking at the result of the signature analyses, through one sample t-tests against zero, we found evidence for a loading of the FaceDonation condition for NPS (W=343, p=0.003, BF_+0_=12.975), for VPS (W=362, p<0.001, BF_+0_=242.022) and also for the guilt signature (t_(28)_=2.289, *p*=0.015, Cohen’s d=0.425, BF_+0_=3.604). The same was the case for the HandDonation condition for NPS (t_(28)_=2.010, *p*=0.027, Cohen’s d=0.373, BF_+0_=2.203), VPS (t_(28)_=1.795, *p*=0.042, Cohen’s d=0.333, BF_+0_=1.548) and the guilt signature (t_(28)_=3.581, *p*<0.001 Cohen’s d=0.665, BF_+0_=53.887) (**Fig. 6**). These results show that both the pain observation network and pain network of our participants covaries with the helping behavior. The same seems to be the case for the guilt network. For the NPS and VPS for the FaceDonation a Shapiro-Wilk’s test rejected the null-hypothesis of a normal distribution, thus a non-parametric Wilcoxon signed rank test vs zero was used.

**Figure 6.**
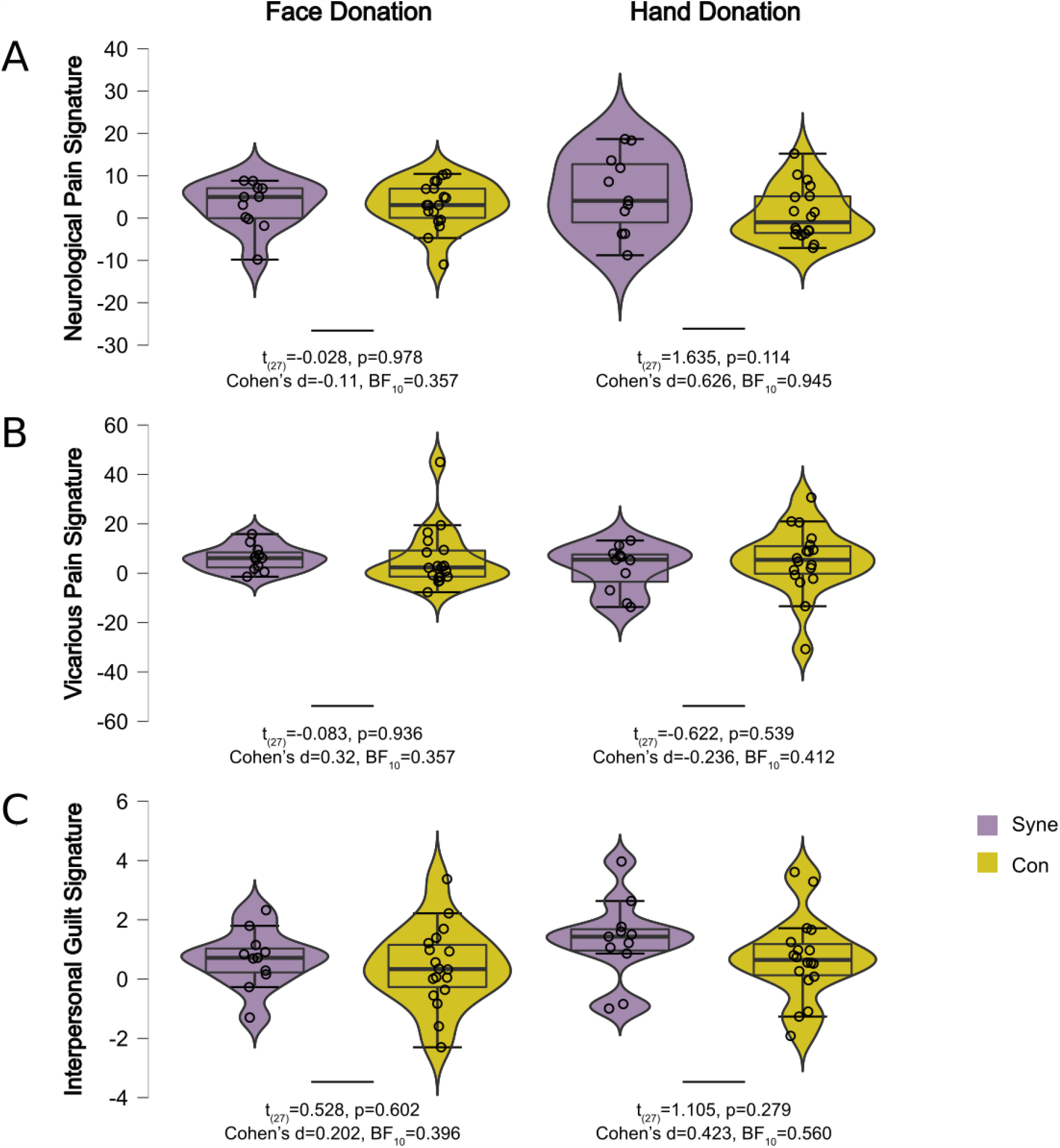
**(A)** Results of the Neurological Pain Signature analyses for the FaceDonation and HandDonation. **(B)** Results of the Vicarious Pain Signature analyses for the FaceDonation and HandDonation. **(C)** Results of the Interpersonal Guilt Signature analyses for the FaceDonation and HandDonation. All results are displayed in arbitrary units. Violin plots represent the distribution, the box-plot within the median and the whisker the quartiles. The BF_10_ and p-values between the violin plots represent the results of the comparison between individuals reporting mirror-pain synesthesia (cyan) experiences and those who did not (yellow).

Comparing the loadings on the three signatures of the FaceDonation and HandDonation between individuals reporting mirror-pain synesthesia experiences and those who did not, did not show any significant differences and there seemed to be evidence of absence of a difference. Comparison mirror-pain self-report synesthesia vs non-mirror-pain reports for FaceDonation: NPS (t_(27)_=-0.028, p=0.978, Cohen’s d=-0.11, BF_10_=0.357), VPS (t_(27)_=0.083, p=0.934, Cohen’s d=0.032, BF_10_=0.357) and for the guilt signature (t_(27)_=0.528, p=0.602, Cohen’s d=0.202, BF_10_=0.396). For HandDonation: NPS (t_(27)_=1.635, p=0.114, Cohen’s d=0.626, BF_10_=0.945), VPS (t_(27)_=-0.622, p=0.539, Cohen’s d=-0.236, BF_10_=0.412) and the guilt signature (t_(27)_=1.105, p=0.279, Cohen’s d=0.423, BF_10_=0.560) (**Fig. 6**).

## Discussion

To shed light onto the contribution of somatic feelings while witnessing the pain of others onto decision-making in costly helping, we contrasted the choices and brain activity of participants that report feeling such somatic feelings (self-reported mirror-pain synesthetes) against those that do not.

Behaviorally, in line with the notion that somatic feelings may contribute to a motivation to help, individuals reporting mirror-pain synesthesia donated more money to reduce the pain of another individual, and their donations increased more steeply as the witnessed pain became more intense. Somewhat surprisingly, this was true whether the pain was perceived from the Face or Hand, although we had expected this to be more strongly the case for the Hand stimuli that were designed to invite viewers to mirror their observation more specifically on their own hand.

Neurally, in addition to finding that brain activity in regions associated with the affective components of pain were associated with donation (including the insula and cingulate for the Face and, at reduced threshold, for the Hand stimuli, in line with previous studies (Hein et al., 2010; Ma et al., 2011; FeldmanHall et al., 2015; Tomova et al. 2016), our results showed that donations were also associated with activity in somatosensory brain regions (SII and, when using a multivariate approach, SI). In addition, as expected, we did find that donations were more tightly associated with activity in the somatosensory cortices (SII) for the self-report synesthetes. This latter finding is in line with previous reports that associate mirror synesthesia with increased somatosensory activation (Blakemore et al., 2005; Osborn and Derbyshire 2010; Grice-Jackson et al., 2017), but extend this finding to prosociality.

In addition, using the neurological pain signature (Wager et al., 2013), developed to quantify the recruitment of neural activity typical of feeling somatic pain on one’s own body, we could confirm that trials with higher donations (as captured by the HandDonation or FaceDonation parametric modulator) were associated with higher recruitment of this somatic pain pattern. Surprisingly, this was true for the Hand and Pain stimuli, without significant differences between them, and was not more strongly the case for self-report synesthetes. This lack of specificity may be due to the holistic nature of the pattern that includes but is not specific to somatosensory brain regions (Wager et al., 2013). That patterns trained to capture the recruitment of pain observation and guilt also overlap with those for costly helping is perhaps less surprising given that several voxelwise studies have associated individual brain regions involved in affective empathy and guilt with prosociality (Hein et al., 2010; Ma et al., 2011; Xu et al., 2014; FeldmanHall et al., 2015; Erlandsson et al., 2016; Tomova et al. 2016) and these more affective regions would be less expected to be associated with self-reported mirror pain synesthesia.

Together, our neuroscientific findings therefore clearly support the notion that the degree to which observers recruit their own pain circuitry, including affective and somatosensory pain components, is indeed associated with their willingness to sacrifice their own money to help others in pain. Our results also support the specific question of how much somatically feeling the pain of others onto our own body contributes to this willingness to help. Self-declared mirror-pain synesthetes, and to a lesser extend responders in the VPQ, do donate more money to alleviate the pain of others in our sample, and have a steeper donation slope (i.e. increase their donations more steeply as the pain of the victim increases). In addition, fMRI analysis revealed that activity in somatosensory cortices is associated with donation, as is the recruitment of patterns associated with somatic pain and activity in SII was more tightly associated with donation for these synesthetes when observing a hand being slapped. The latter suggests that if participants do somatically mirror the pain of others onto their own body, this leads to an increased motivation that is also more dependent on activity in their secondary somatosensory cortices.

Our study also has certain limitations that could inspire future studies. Firstly, we used self-reports of mirror pain experiences outside of the lab as our way to identify who is a mirror-pain synesthetes. Philosophically, it is such subjective nociception in real life that is thought to motivate helping. However, future studies may wish to probe how much mirror pain participants feel on every trial within the experiment, and examine if variance in mirroring across trials in which similar levels of pain are observed can account for unique variance in helping. That the VPQ for instance does not lead to the exact same classification into mirror pain-synesthetes and controls (see Table 1) -- although it’s classification also leads to significant differences in donation – reinforces the opportunity for more fine grained analysis of the subjective experience of mirror pain and its association with helping. Second, our neuroimaging findings only show significant correlations between brain activity (or multivariate patterns thereof) in somatosensory regions and donations, and cannot prove that such associations are causal in nature. In the past, we have shown that altering brain activity in SI non-invasively using TMS in participants can alter helping, specifically by altering how tightly participants tailor their helping to the needs of the target individual (Gallo et al., 2018). Using similar methodologies in participants reporting mirror pain synesthesia and measuring the effect on subjective feelings and helping will be key to a tighter understanding of the contribution of somatosensory cortices, the difference between SI and SII’s contribution, and the motivation to help. Finally, we only tested female participants in our paradigm. This was a decision we made based on extensive literature showing that synesthesia is by far more common in women (Baron-Cohen et al., 1996; Calkins, 1895; Domino, 1989; Ramachandran and Hubbard, 2001) even though there is some evidence that the difference is partly biased due to the fact the female participants are reacting more to experiment calls (Simner and Carmichael, 2015). We were also aware that Gallo et al. (2018) using the same paradigm did not find any differences in donation between females and males. Future studies may however attempt to recruit more male mirror-touch synesthetes to explore if there might be sex-differences in the circuitry motivating helping, and in the contribution of somatosensory regions in particular. That preclinical studies have suggested substantial differences in the biological basis of nociception across male and female rodents is an intriguing reminder for the need of being mindful of the potential for sex-differences in pain-related phenomena (Mogil, 2020).

We could not find out more on why participants engaged to help through scales of questionnaires such as the IRI, measuring empathy, or MAS, measuring attitude towards money, since none of these measures correlated with the donation made. When taking into account research showing that empathy is determined both by context dependency and automaticity (Zaki, 2014) and that the balance between ability and propensity to empathize can play a key role in different populations (Keysers and Gazzola, 2014), this becomes less surprising. Both the measures we used are trait and not state measures and not developed for use under conflictual contexts.

## Acknowledgements

The research was funded by the Netherlands Organization for Scientific Research (VIDI 452– 14–015 to V.G.). We thank M. Spezio for discussion during the development of the paradigm.

## Author contributions

V.G, C.K., K.I. and S.G. designed the study. K.I. and S.G. set up the experiment. S.G. was the confederate and data collection was performed by K.I. The data analysis was performed by K.I. under the supervision of V.G. and C.K. and S.G. contributed to the first round of analyses. K.I., V.G. and C.K. wrote the manuscript. All authors approved the final version of the manuscript for submission.

## Data availability

Data are made available on OSF at https://osf.io/d3vyq/

## Competing Interests

The authors declare that there is no conflict of interest.

## SUPPLEMENTARY INFORMATION S1

### Vicarious Pain Questionnaire

#### Method

The VPQ (Grice-Jackson et al., 2017) consists of 16 video clips of 10 seconds each depicting painful situations (e.g. injections and sporting injuries). After watching each video participants were asked if they felt any pain on their own body. In case they gave a positive response they were then asked additional questions: to rate the intensity (using a scale from 1 to 10), to report the location of the pain as it was felt (localized in same location as observed, localized in another location, a non localized general sensation), to choose any word from a list of pain adjectives that matched their vicarious pain experience. We followed the same two-step cluster analysis approach as in Grice-Jackson et al., 2017 which resulted in three different groups: affective generalized group A/G, sensory localized group S/L and non-responders. The tool was administered via LimeSurvey platform (www.limesurvey.org). Three participants did not fill it in (one that had reported mirror pain synesthesia experiences and two that had not).

#### Results

Out of the 31 participants that were included in our behavioral analyses, three participants did not complete the VPQ. Following the classification method of Grice-Jackson et al. (2017), of the 28 participants that did complete the VPQ, the distribution differed based on whether participants report mirror touch synesthesia or not (χ^2^(df=2)=7.032, *p*=0.03, BF_10_=5.283), with the likelihood to be classified as sensory/localizer 8 times higher in participants that reported mirror touch synesthesia than in those that do not (see Table S1). However, not all participants that self-report mirror touch synesthesia do qualify as responders.

**Table S1:** VPQ classification as a function of self-report. Each cell contains the number and proportion of participants falling into a specific VPQ classification (column) as a function of whether they do (top) or do not (bottom) self-report mirror pain synesthesia experiences in everyday life. A Chi-Square test on the contingency table confirms a significant difference in VPQ distribution based on self-report status, with self-reported mirror pain synesthetes having a higher proportion of sensory localizers and a lower proportion of non-responders (χ^2^(df=2)=7.032, *p*=0.03, BF_10_=5.283).

|  | Sensory Localizers | Affective Generalizers | Non Responders | Missing VPQ | Total |
| --- | --- | --- | --- | --- | --- |
| Self Reported Mirror Pain Synesthesia | 6/12=50% | 1/12=8% | 5/12=42% | 1 | 13 |
| No Self-Report of Mirror Pain Synesthesia | 1/16=6% | 2/16=12% | 13/16=81% | 2 | 18 |

**Table S2:**
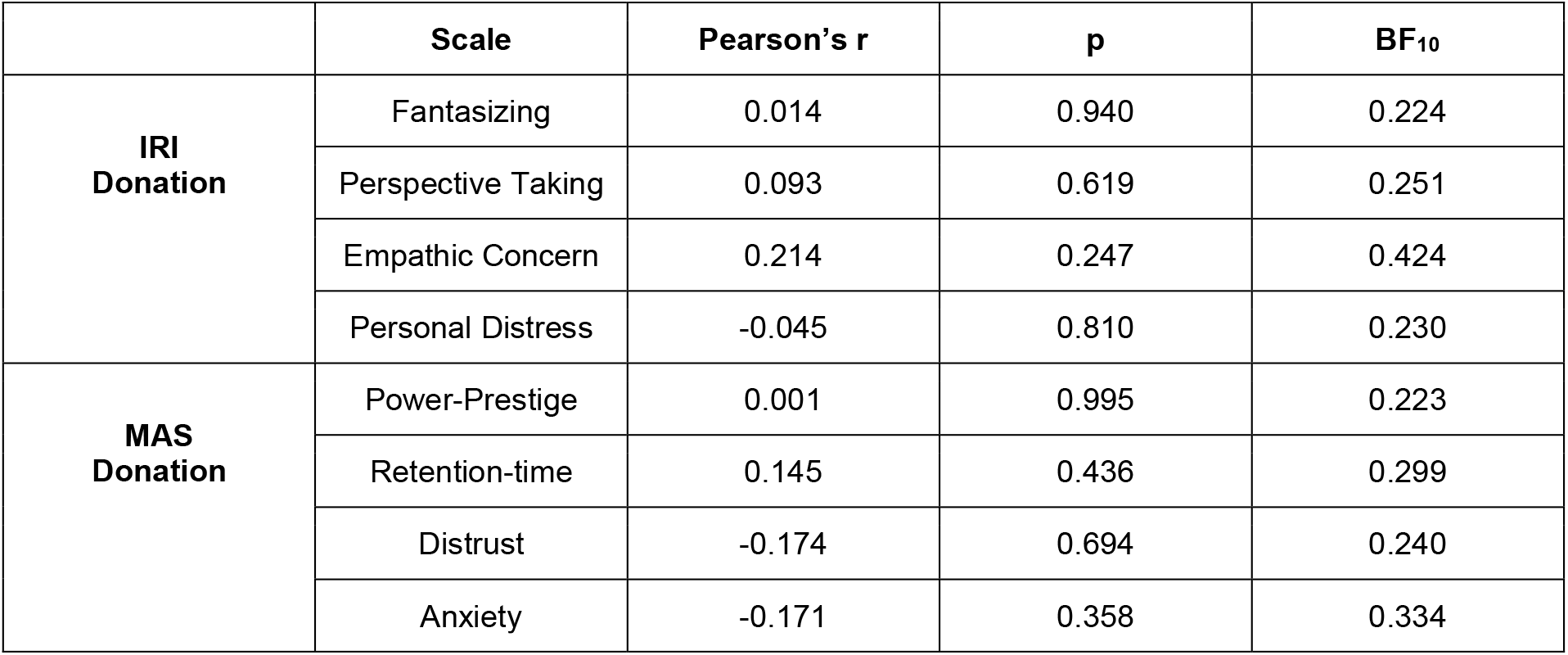
Correlations between IRI, MAS and the average donation that participants made. The table summarizes the correlations for the average donation that participants made for the Face and Hand conditions together and the subscales of the IRI (Davis and Association, 1980) (Fantasizing, Perspective Taking, Empathic Concern and Personal Distress) and MAS (Yamauchi and Templer, 1982) (Power-Prestige, Retention-time, Distrust and Anxiety). None of these correlations were significant (all p>0.05) and all BFs<0.424 suggesting evidence for absence of an effect.

## SUPPLEMENTARY INFORMATION S2

### Univariate GLM analyses

**Figure S1.**
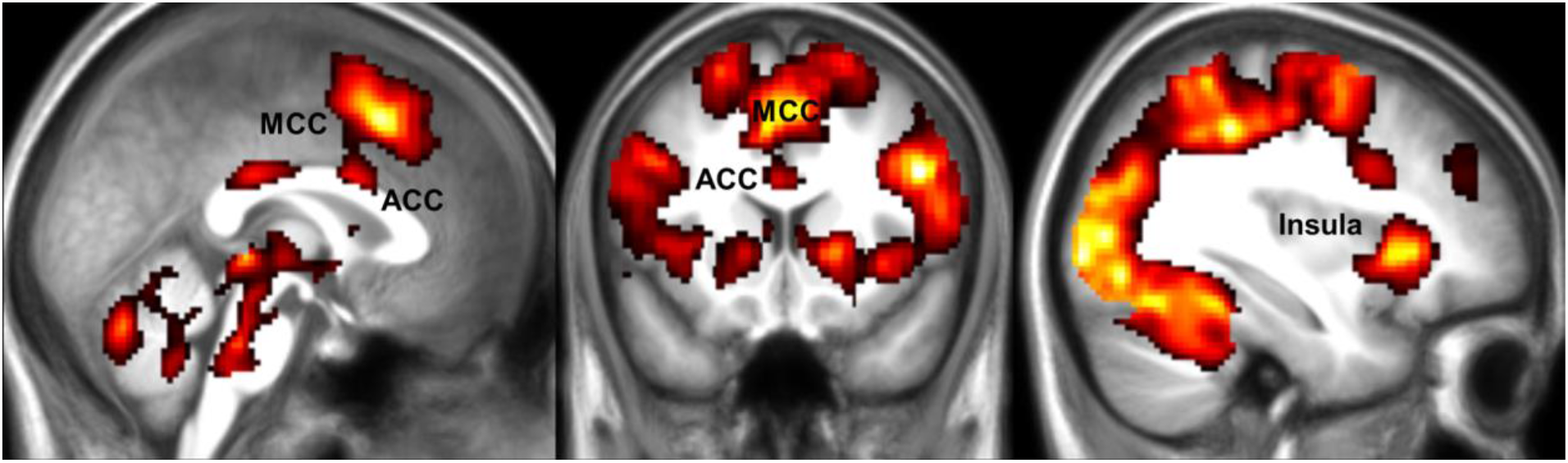
Main effect of video 1. Results of the main effects of Face and Hand conditions together, indicating voxels where BOLD signals during the pain observation are increased. Results are FWE cluster-corrected at p<0.05 (p<0.001, k=FWEc=389 voxels, cFWE 3.34<t<13).

**Table S3:**
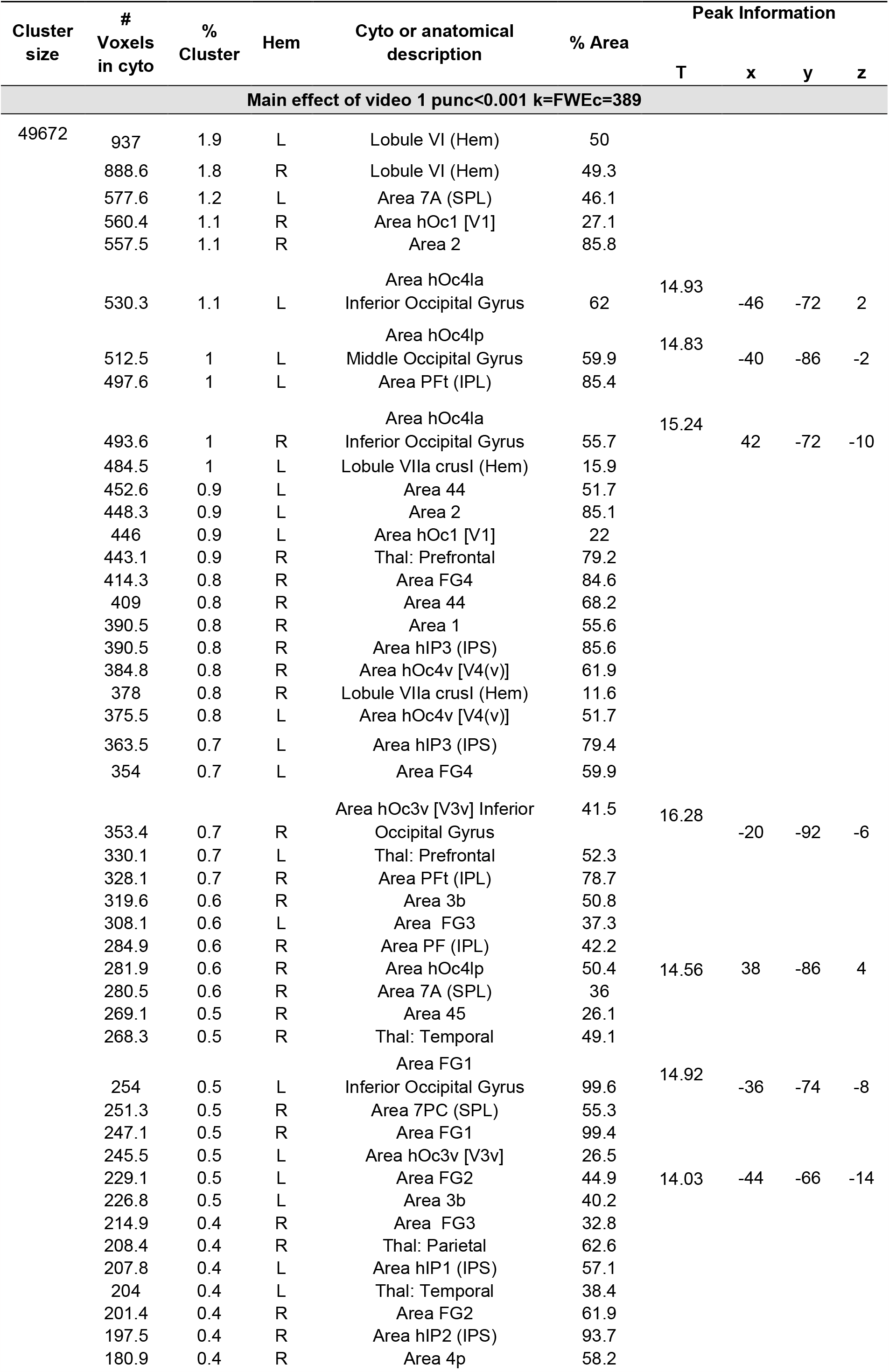
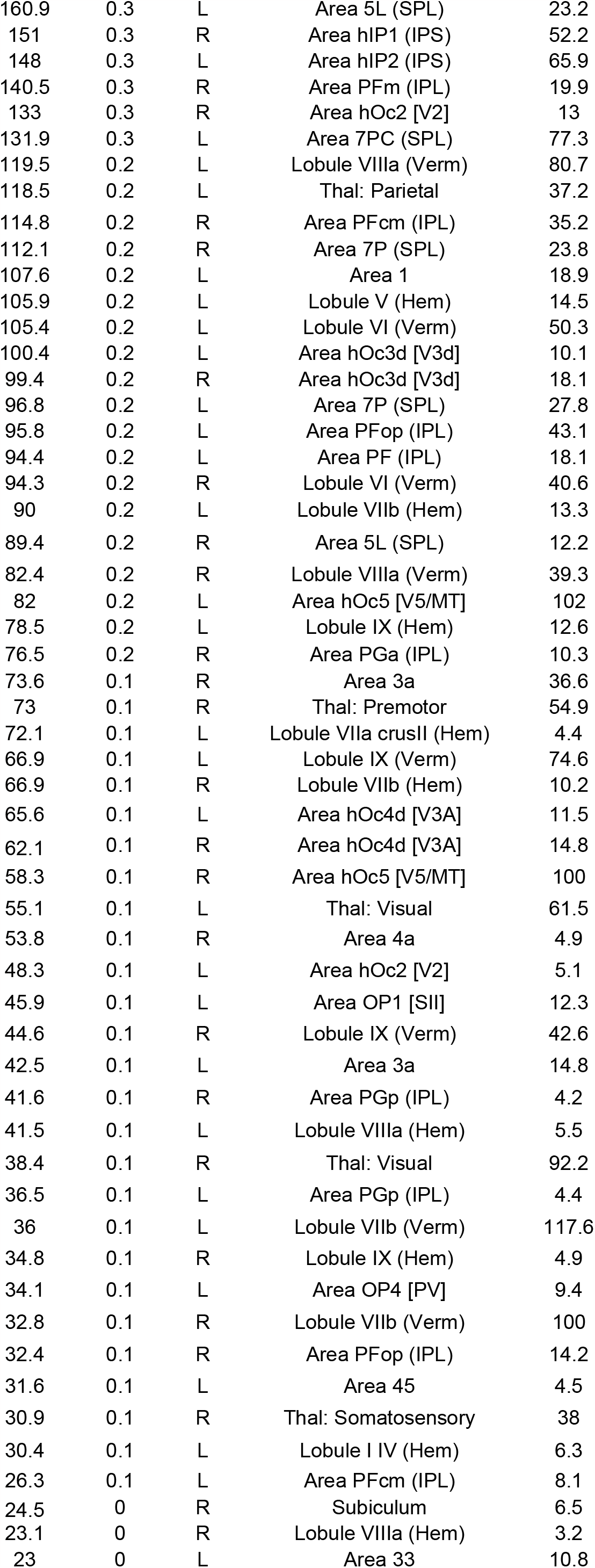
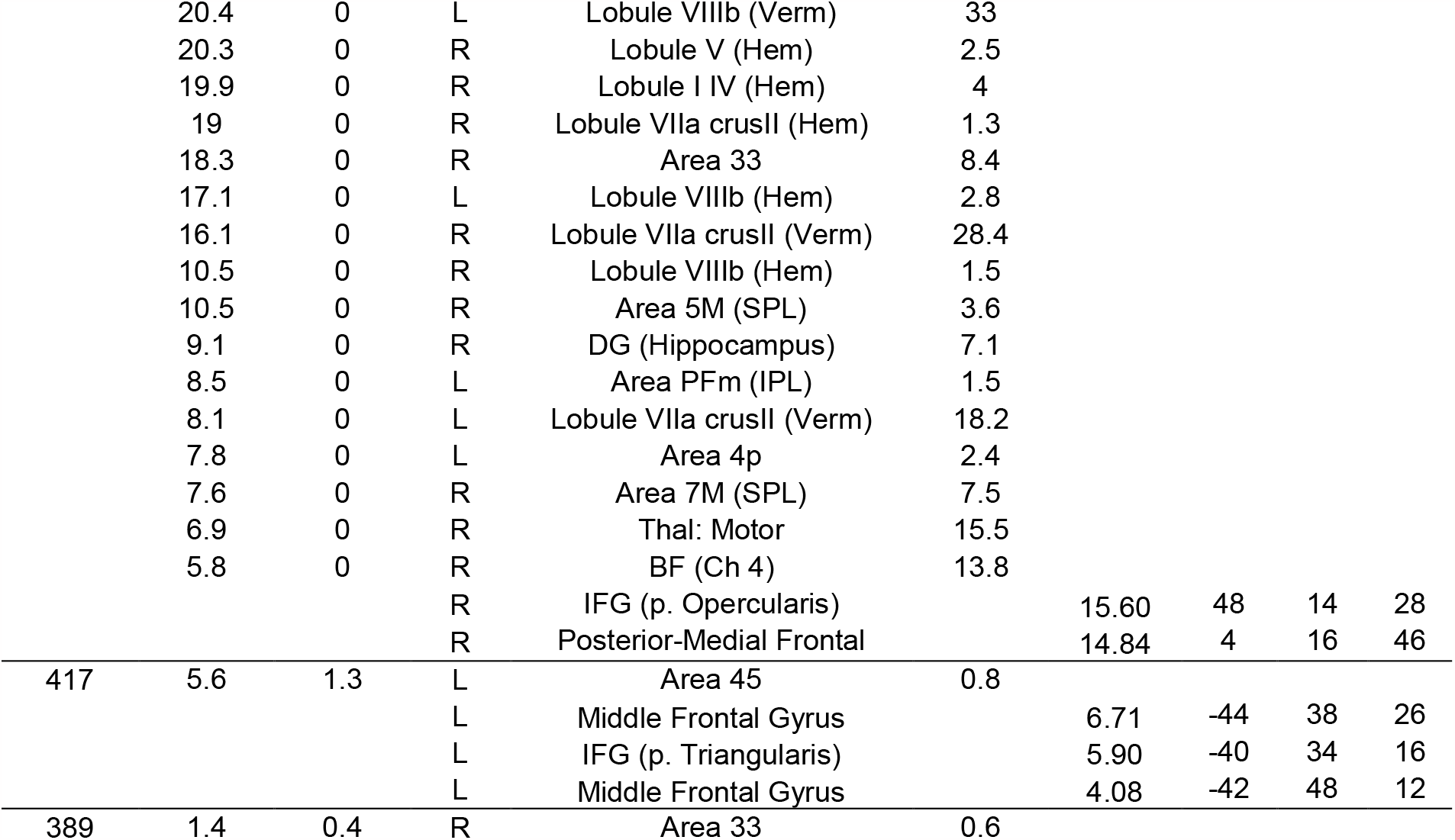
Results of the voxelwise analysis. Brain activations for the main effect of video 1. Regions were labeled using SPM Anatomy Toolbox. From left to right: the cluster size in number of voxels, the number of voxels falling in a cyto-architectonic area, the percentage of the cluster that falls in the cyto-architectonic area, the hemisphere (L=left; R=right), the name of the cyto-architectonic area when available or the anatomical description, the percentage of the area that is activated by the cluster, the t values of the peaks associated with the cluster followed by their MNI coordinates in mm.

**Figure S2.**
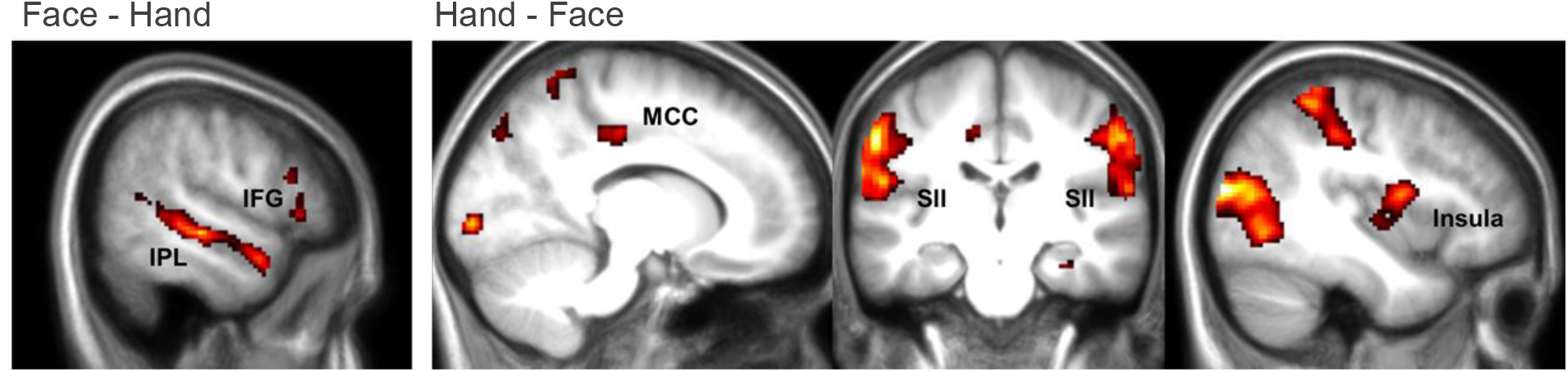
Effect of stimulus type (Face vs Hand). Comparison between the face and hand conditions for the first video pain observation. Results are FWE cluster-corrected at p<0.05 (p_unc_<0.001, k=FWEc=145 voxels for the face and k=FWEc=125 voxels for the hand). For this analysis we constructed a different GLM, same to the one described at the *GLM analysis* methods section, without any parametric modulators.

**Table S4:**
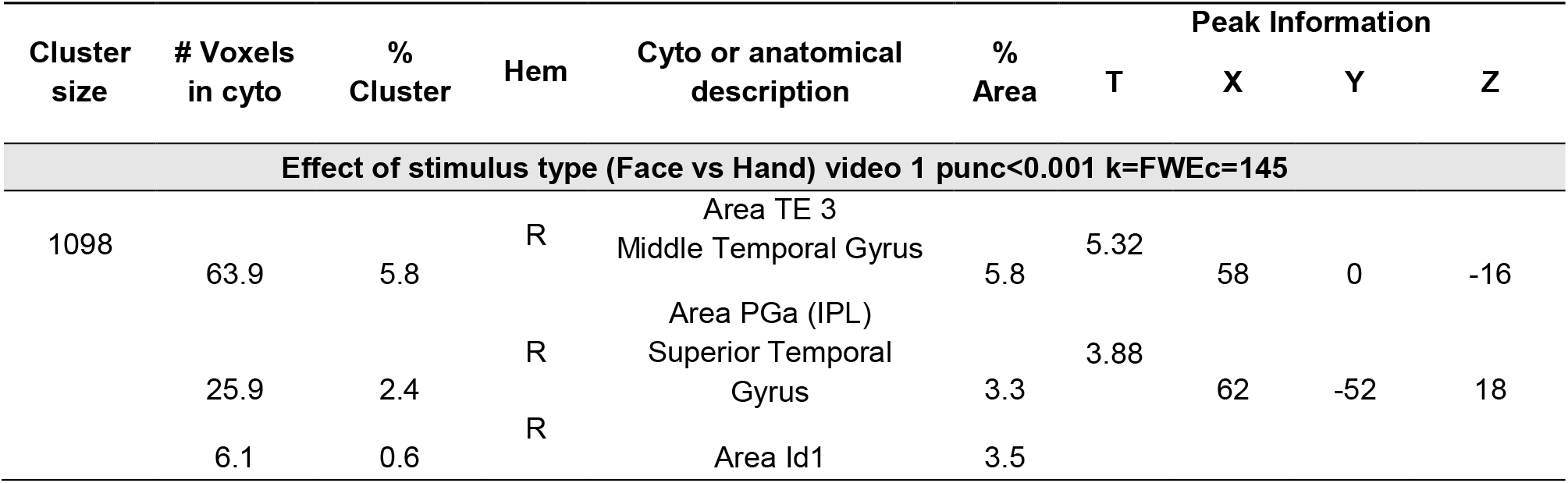
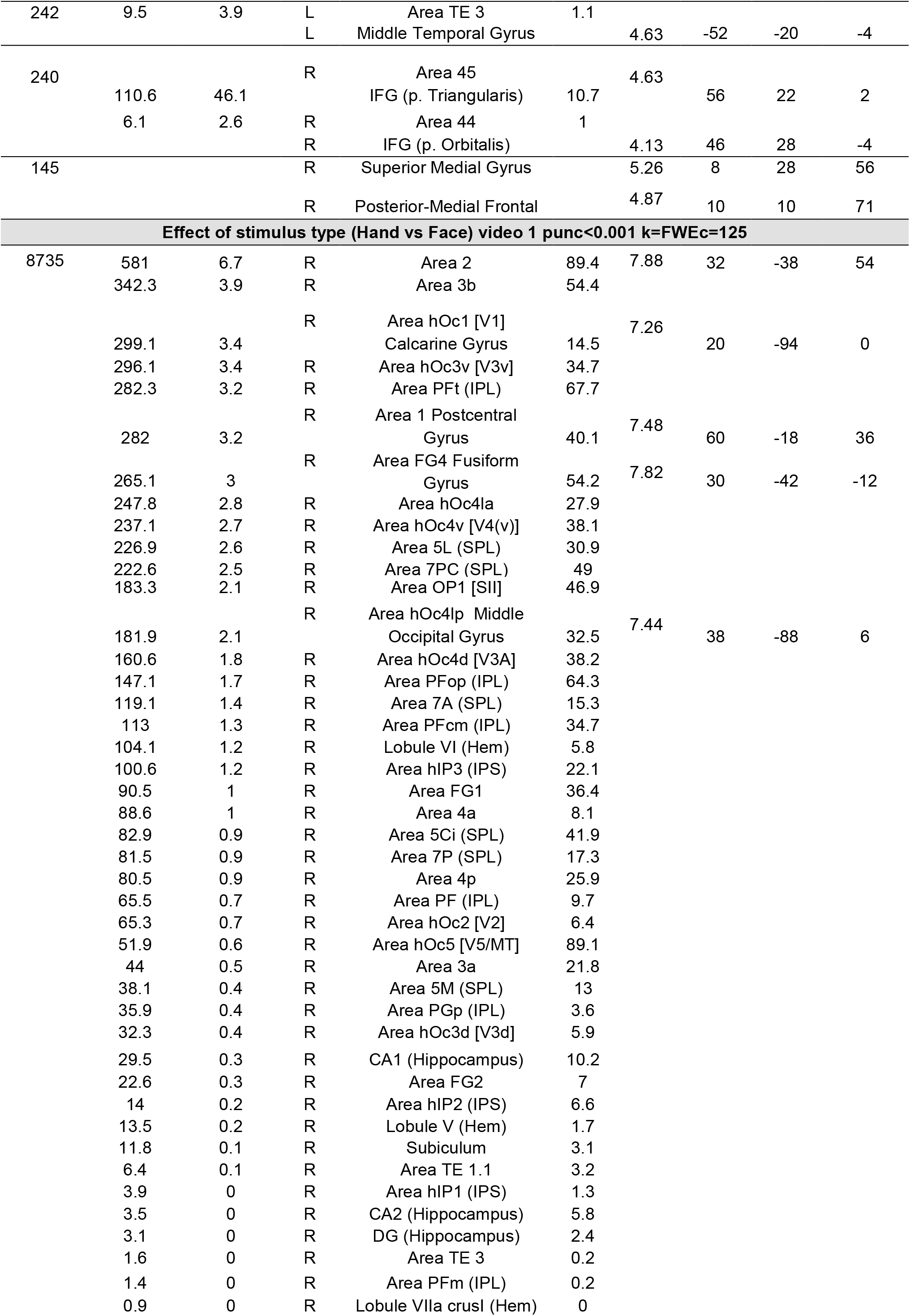
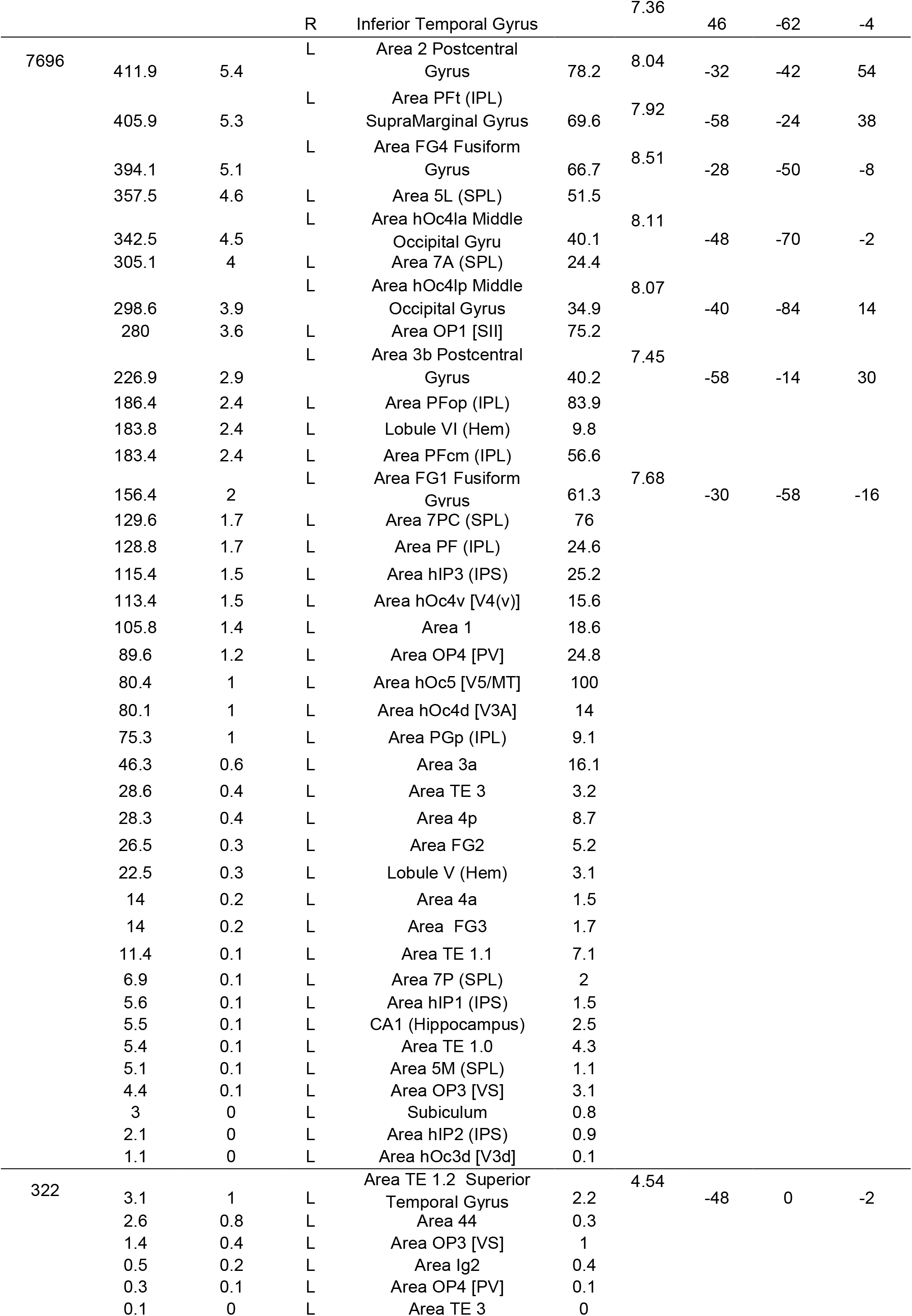
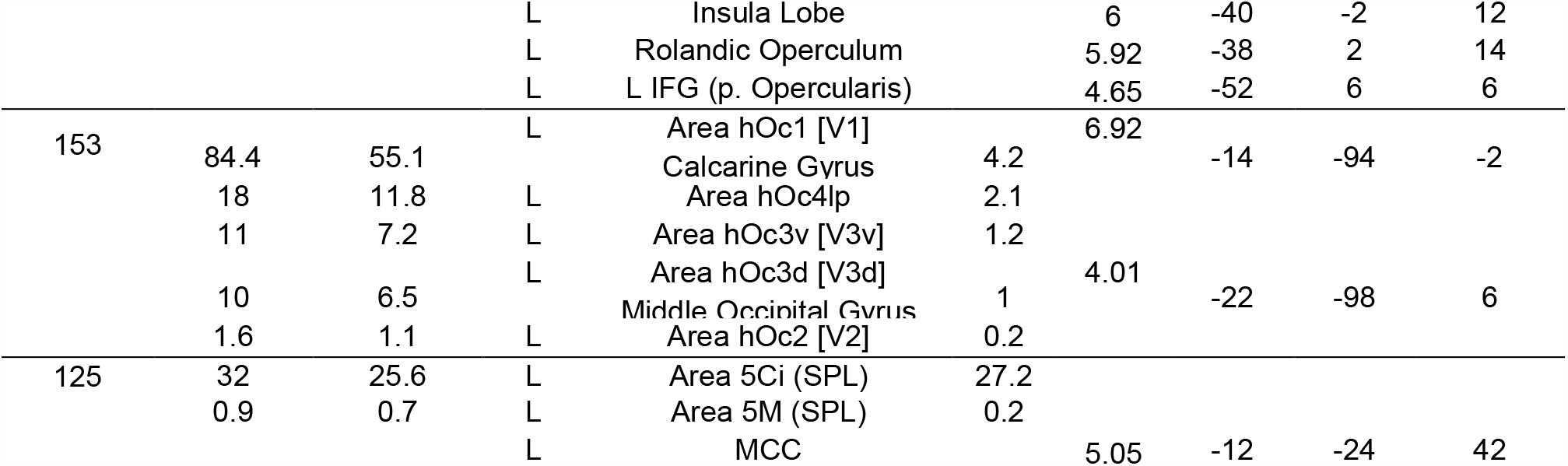
Results of the voxelwise analysis. Brain activations for the effect of stimulus type (Face vs Hand). Regions were labeled using SPM Anatomy Toolbox. From left to right: the cluster size in number of voxels, the number of voxels falling in a cyto-architectonic area, the percentage of the cluster that falls in the cyto-architectonic area, the hemisphere (L=left; R=right), the name of the cyto-architectonic area when available or the anatomical description, the percentage of the area that is activated by the cluster, the t values of the peaks associated with the cluster followed by their MNI coordinates in mm.

**Figure S3.**
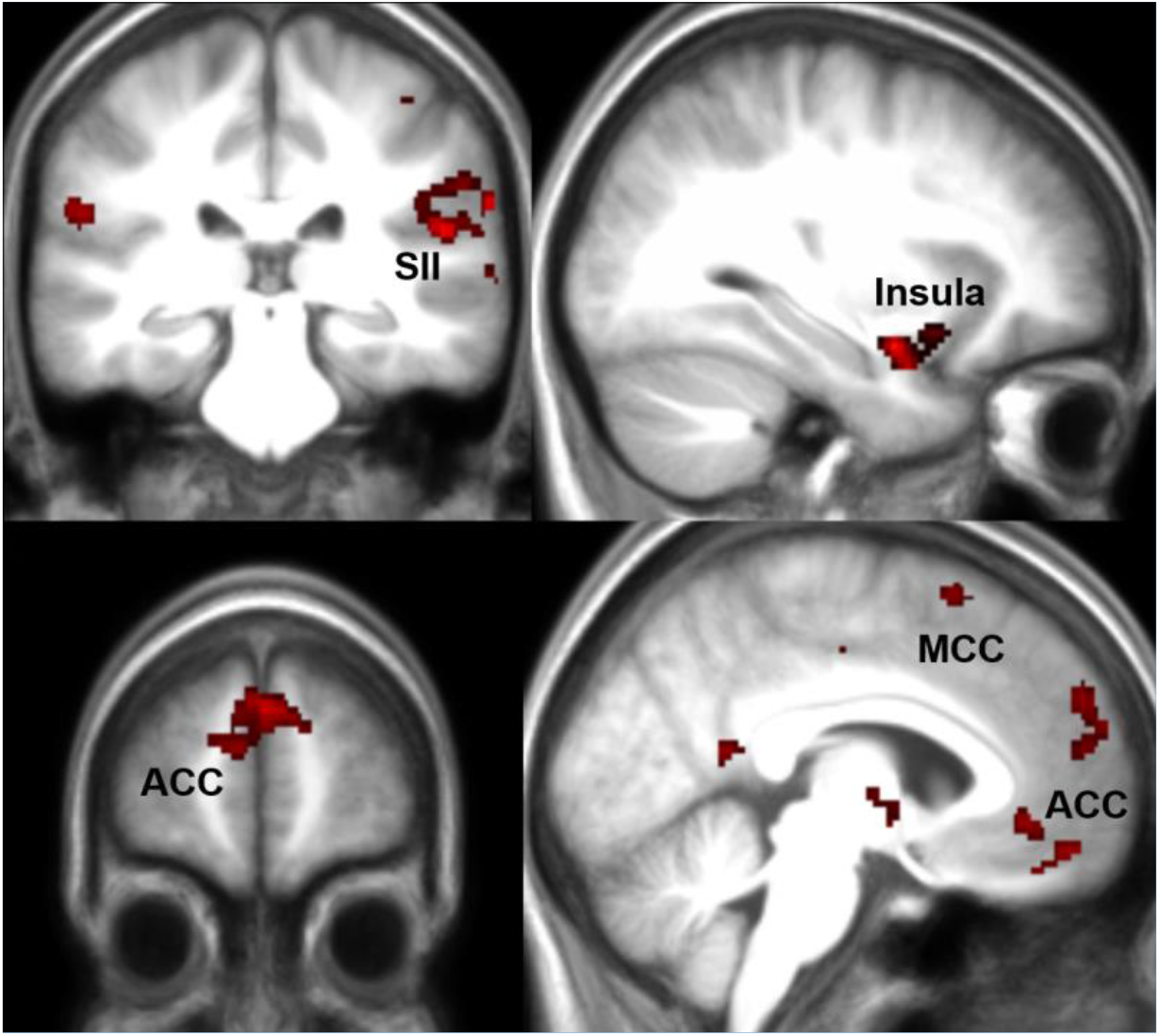
HandDonation parametric modulator at reduced threshold. Results of a linear regression on the parametric modulator for the first video and trial-by-trial donation in the Hand condition. This identifies voxels with signals that increase for higher donation. Results are shown at uncorrected p<0.01, 2.4<t<8.

**Table S5:**
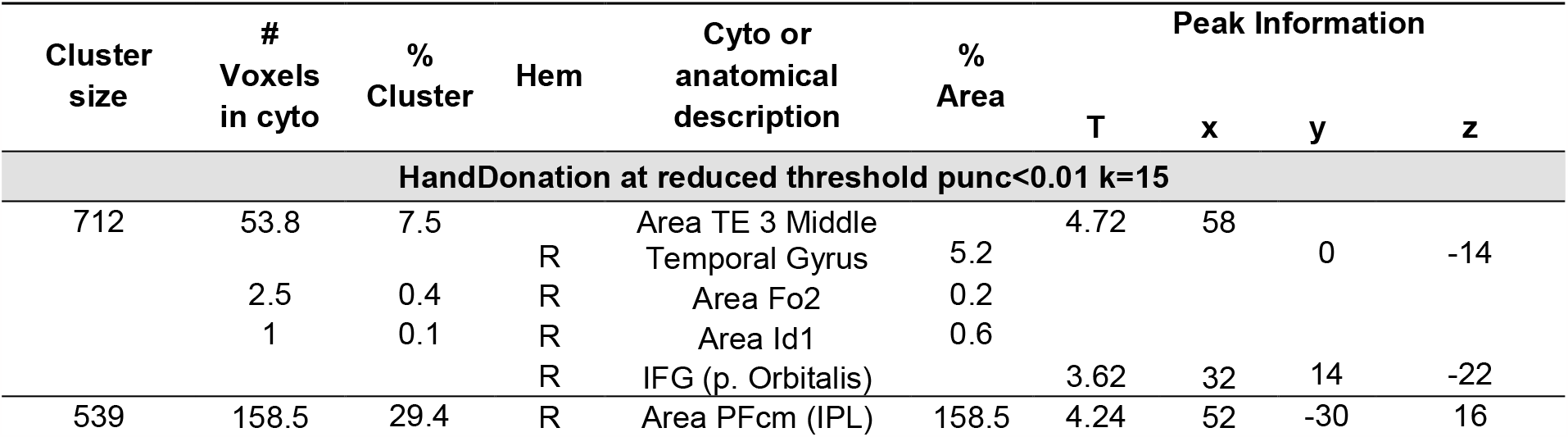
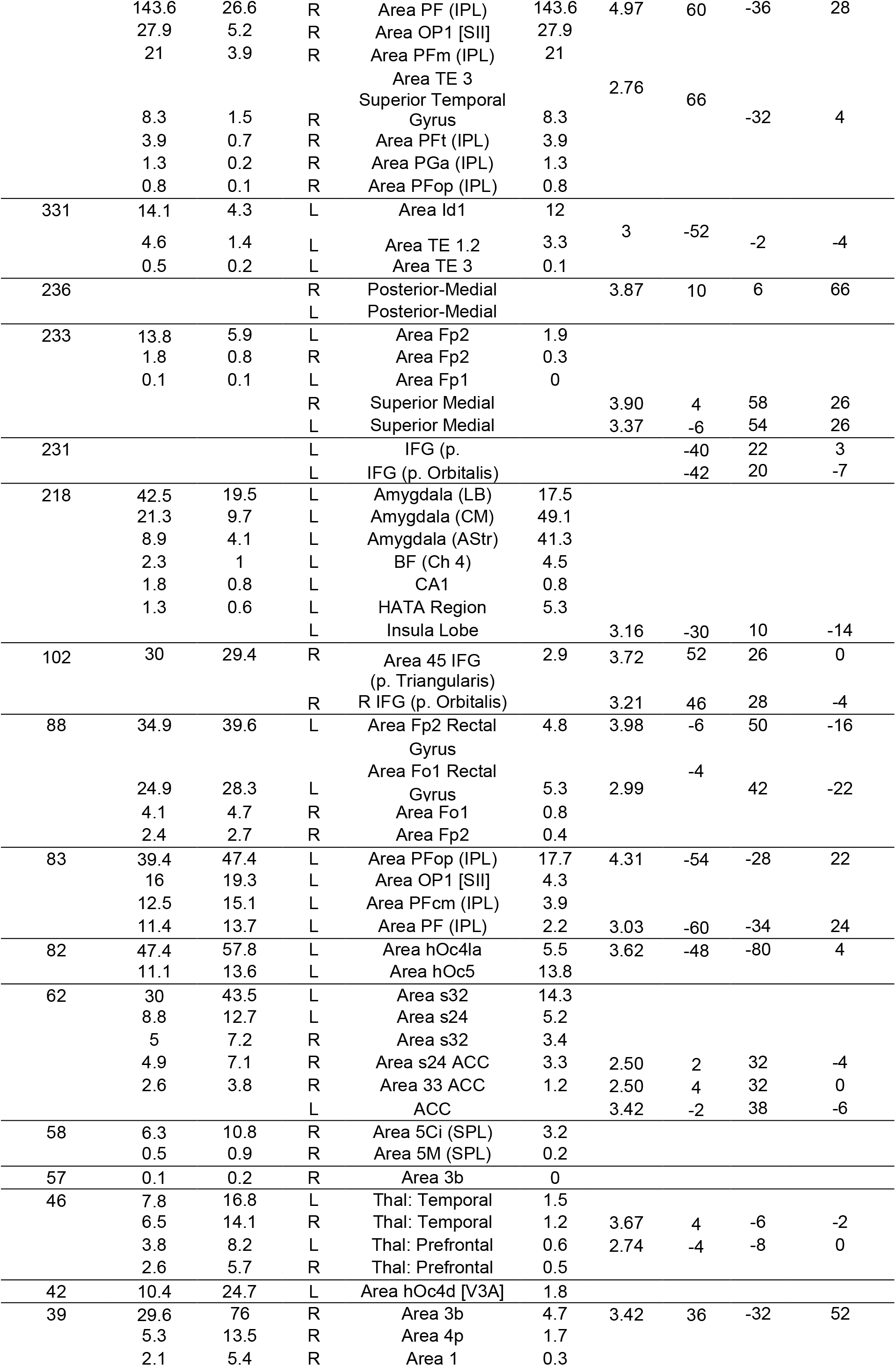
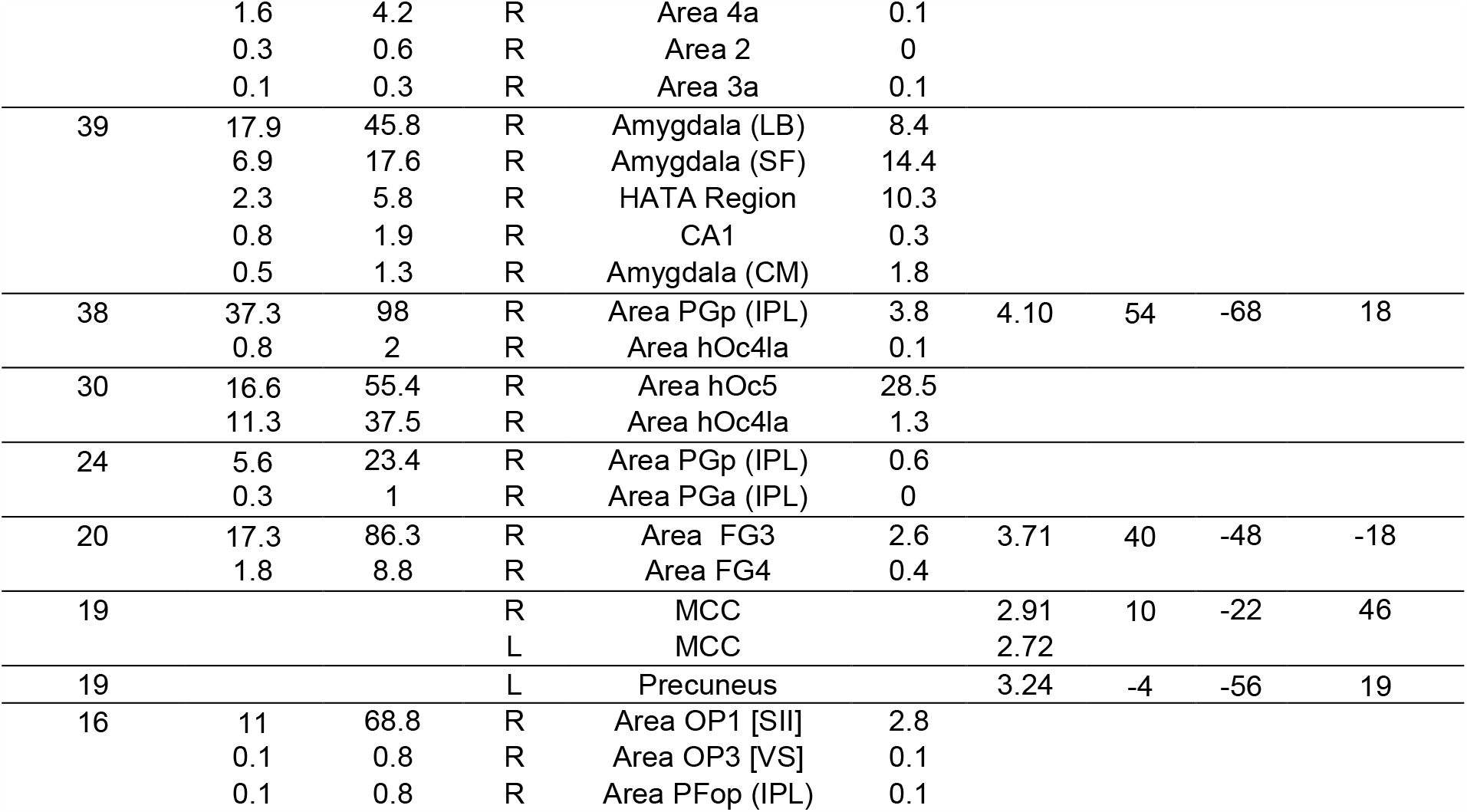
Results of the voxelwise analysis. Brain activations for the HandDonation parametric modulator for all participants together at reduced threshold. Regions were labeled using SPM Anatomy Toolbox. From left to right: the cluster size in number of voxels, the number of voxels falling in a cyto-architectonic area, the percentage of the cluster that falls in the cyto-architectonic area, the hemisphere (L=left; R=right), the name of the cyto-architectonic area when available or the anatomical description, the percentage of the area that is activated by the cluster, the t values of the peaks associated with the cluster followed by their MNI coordinates in mm.

| Cluster size | # Voxels in cyto | % Cluster | Hem | Cyto or anatomical description | % Area | Peak Information |  |  |  |
| --- | --- | --- | --- | --- | --- | --- | --- | --- | --- |
|  |  |  |  |  |  | T | x | y | z |
| HandDonation at reduced threshold punc<0.01 k=15 |  |  |  |  |  |  |  |  |  |
| 712 | 53.8 | 7.5 |  | Area TE 3 Middle |  | 4.72 | 58 |  |  |
|  |  |  | R | Temporal Gyrus | 5.2 |  |  | 0 | -14 |
|  | 2.5 | 0.4 | R | Area Fo2 | 0.2 |  |  |  |  |
|  | 1 | 0.1 | R | Area Id1 | 0.6 |  |  |  |  |
|  |  |  | R | IFG (p. Orbitalis) |  | 3.62 | 32 | 14 | -22 |
| 539 | 158.5 | 29.4 | R | Area PFcm (IPL) | 158.5 | 4.24 | 52 | -30 | 16 |
|  | 143.6 | 26.6 | R | Area PF (IPL) | 143.6 | 4.97 | 60 | -36 | 28 |
|  | 27.9 | 5.2 | R | Area OP1 [SII] | 27.9 |  |  |  |  |
|  | 21 | 3.9 | R | Area PFm (IPL) | 21 |  |  |  |  |
|  |  |  |  | Area TE 3 |  | 2.76 |  |  |  |
|  |  |  |  | Superior Temporal |  |  | 66 |  |  |
|  | 8.3 | 1.5 | R | Gyrus | 8.3 |  |  | -32 | 4 |
|  | 3.9 | 0.7 | R | Area PFt (IPL) | 3.9 |  |  |  |  |
|  | 1.3 | 0.2 | R | Area PGa (IPL) | 1.3 |  |  |  |  |
|  | 0.8 | 0.1 | R | Area PFop (IPL) | 0.8 |  |  |  |  |
| 331 | 14.1 | 4.3 | L | Area Id1 | 12 |  |  |  |  |
|  | 4.6 | 1.4 | L | Area TE 1.2 | 3.3 | 3 | -52 | -2 | -4 |
|  | 0.5 | 0.2 | L | Area TE 3 | 0.1 |  |  |  |  |
| 236 |  |  | R | Posterior-Medial |  | 3.87 | 10 | 6 | 66 |
|  |  |  | L | Posterior-Medial |  |  |  |  |  |
| 233 | 13.8 | 5.9 | L | Area Fp2 | 1.9 |  |  |  |  |
|  | 1.8 | 0.8 | R | Area Fp2 | 0.3 |  |  |  |  |
|  | 0.1 | 0.1 | L | Area Fp1 | 0 |  |  |  |  |
|  |  |  | R | Superior Medial |  | 3.90 | 4 | 58 | 26 |
|  |  |  | L | Superior Medial |  | 3.37 | -6 | 54 | 26 |
| 231 |  |  | L | IFG (p. |  |  | -40 | 22 | 3 |
|  |  |  | L | IFG (p. Orbitalis) |  |  | -42 | 20 | -7 |
| 218 | 42.5 | 19.5 | L | Amygdala (LB) | 17.5 |  |  |  |  |
|  | 21.3 | 9.7 | L | Amygdala (CM) | 49.1 |  |  |  |  |
|  | 8.9 | 4.1 | L | Amygdala (AStr) | 41.3 |  |  |  |  |
|  | 2.3 | 1 | L | BF (Ch 4) | 4.5 |  |  |  |  |
|  | 1.8 | 0.8 | L | CA1 | 0.8 |  |  |  |  |
|  | 1.3 | 0.6 | L | HATA Region | 5.3 |  |  |  |  |
|  |  |  | L | Insula Lobe |  | 3.16 | -30 | 10 | -14 |
| 102 | 30 | 29.4 | R | Area 45 IFG | 2.9 | 3.72 | 52 | 26 | 0 |
|  |  |  |  | (p. Triangularis) |  |  |  |  |  |
|  |  |  | R | R IFG (p. Orbitalis) |  | 3.21 | 46 | 28 | -4 |
| 88 | 34.9 | 39.6 | L | Area Fp2 Rectal | 4.8 | 3.98 | -6 | 50 | -16 |
|  |  |  |  | Gyrus |  |  |  |  |  |
|  |  |  |  | Area Fo1 Rectal |  |  | -4 |  |  |
|  | 24.9 | 28.3 | L | Gyrus | 5.3 | 2.99 |  | 42 | -22 |
|  | 4.1 | 4.7 | R | Area Fo1 | 0.8 |  |  |  |  |
|  | 2.4 | 2.7 | R | Area Fp2 | 0.4 |  |  |  |  |
| 83 | 39.4 | 47.4 | L | Area PFop (IPL) | 17.7 | 4.31 | -54 | -28 | 22 |
|  | 16 | 19.3 | L | Area OP1 [SII] | 4.3 |  |  |  |  |
|  | 12.5 | 15.1 | L | Area PFcm (IPL) | 3.9 |  |  |  |  |
|  | 11.4 | 13.7 | L | Area PF (IPL) | 2.2 | 3.03 | -60 | -34 | 24 |
| 82 | 47.4 | 57.8 | L | Area hOc4la | 5.5 | 3.62 | -48 | -80 | 4 |
|  | 11.1 | 13.6 | L | Area hOc5 | 13.8 |  |  |  |  |
| 62 | 30 | 43.5 | L | Area s32 | 14.3 |  |  |  |  |
|  | 8.8 | 12.7 | L | Area s24 | 5.2 |  |  |  |  |
|  | 5 | 7.2 | R | Area s32 | 3.4 |  |  |  |  |
|  | 4.9 | 7.1 | R | Area s24 ACC | 3.3 | 2.50 | 2 | 32 | -4 |
|  | 2.6 | 3.8 | R | Area 33 ACC | 1.2 | 2.50 | 4 | 32 | 0 |
|  |  |  | L | ACC |  | 3.42 | -2 | 38 | -6 |
| 58 | 6.3 | 10.8 | R | Area 5Ci (SPL) | 3.2 |  |  |  |  |
|  | 0.5 | 0.9 | R | Area 5M (SPL) | 0.2 |  |  |  |  |
| 57 | 0.1 | 0.2 | R | Area 3b | 0 |  |  |  |  |
| 46 | 7.8 | 16.8 | L | Thal: Temporal | 1.5 |  |  |  |  |
|  | 6.5 | 14.1 | R | Thal: Temporal | 1.2 | 3.67 | 4 | -6 | -2 |
|  | 3.8 | 8.2 | L | Thal: Prefrontal | 0.6 | 2.74 | -4 | -8 | 0 |
|  | 2.6 | 5.7 | R | Thal: Prefrontal | 0.5 |  |  |  |  |
| 42 | 10.4 | 24.7 | L | Area hOc4d [V3A] | 1.8 |  |  |  |  |
| 39 | 29.6 | 76 | R | Area 3b | 4.7 | 3.42 | 36 | -32 | 52 |
|  | 5.3 | 13.5 | R | Area 4p | 1.7 |  |  |  |  |
|  | 2.1 | 5.4 | R | Area 1 | 0.3 |  |  |  |  |
|  | 1.6 | 4.2 | R | Area 4a | 0.1 |  |  |  |  |
|  | 0.3 | 0.6 | R | Area 2 | 0 |  |  |  |  |
|  | 0.1 | 0.3 | R | Area 3a | 0.1 |  |  |  |  |
| 39 | 17.9 | 45.8 | R | Amygdala (LB) | 8.4 |  |  |  |  |
|  | 6.9 | 17.6 | R | Amygdala (SF) | 14.4 |  |  |  |  |
|  | 2.3 | 5.8 | R | HATA Region | 10.3 |  |  |  |  |
|  | 0.8 | 1.9 | R | CA1 | 0.3 |  |  |  |  |
|  | 0.5 | 1.3 | R | Amygdala (CM) | 1.8 |  |  |  |  |
| 38 | 37.3 | 98 | R | Area PGp (IPL) | 3.8 | 4.10 | 54 | -68 | 18 |
|  | 0.8 | 2 | R | Area hOc4la | 0.1 |  |  |  |  |
| 30 | 16.6 | 55.4 | R | Area hOc5 | 28.5 |  |  |  |  |
|  | 11.3 | 37.5 | R | Area hOc4la | 1.3 |  |  |  |  |
| 24 | 5.6 | 23.4 | R | Area PGp (IPL) | 0.6 |  |  |  |  |
|  | 0.3 | 1 | R | Area PGa (IPL) | 0 |  |  |  |  |
| 20 | 17.3 | 86.3 | R | Area FG3 | 2.6 | 3.71 | 40 | -48 | -18 |
|  | 1.8 | 8.8 | R | Area FG4 | 0.4 |  |  |  |  |
| 19 |  |  | R | MCC |  | 2.91 | 10 | -22 | 46 |
|  |  |  | L | MCC |  | 2.72 |  |  |  |
| 19 |  |  | L | Precuneus |  | 3.24 | -4 | -56 | 19 |
| 16 | 11 | 68.8 | R | Area OP1 [SII] | 2.8 |  |  |  |  |
|  | 0.1 | 0.8 | R | Area OP3 [VS] | 0.1 |  |  |  |  |
|  | 0.1 | 0.8 | R | Area PFop (IPL) | 0.1 |  |  |  |  |

## SUPPLEMENTARY INFORMATION S3

### Multivariate fMRI analysis

#### Methods

For each participant, we performed a general linear model that estimated a separate parameter estimate for the activity during movie 1 for each level of donation (0-6) that participants made at least two times during the experiment. In case a level of donation occurred lust once, it was added in a regressor of no interest and was not analyzed further. Using the anatomy toolbox, we then created an ROI containing all voxels attributed to SI according to the maximum probability maps (i.e. including bilateral BA3a, 3b, 1 and 2), and transformed this mask into the space of the parameter estimates using imagecalc. Using matlab, we then loaded for each subject the parameter estimate images for each level of donation, and only included voxels that fell within our SI mask. Next, we performed a weighted leave-one-subject-out cross-validated partial least square regression. For each of the 29 participants, we kept one subject out, and used the function plsregress in matlab to determine the linear combination of voxels that best predicts donation in the remaining participants. Because some parameter estimates derived from only two trials, and others from as many as 22 trials, we weighted the regression by replicating each parameter estimate image in the training and testing set by the number of trials that went into it. We then used this optimal linear combination to predict the donation of the left-out participant, and quantified the accuracy of the prediction as the correlation between predicted and actual donations. We used Kendall’s Tau as the measure of correlation because it is less susceptible to outliers as a parametric correlation. For the pls-regression, results are shown for using 10 components, based on the elbow method of explained variance including the entire dataset, but results are stable over a range of 8-20 components. We also performed a PCR by first performing a principle component analysis on all the voxel parameter estimates, and then using the first 10 components to perform leave one out regressions to predict donation. This also led to above chance estimates, but in the paper we only report the partial least square regression approach.

#### Results

To explore if SI (i.e. BA3a,3b,1,2) contains information about donation also for the hand trials, for which we failed to find significant evidence at the univariate level that survives correction, we performed a multivariate analysis. Specifically, we trained a weighted partial least-square regression using the data from all but one participant to estimate donation based on a linear combination of the parameter estimates in each voxel in SI, and then used this linear combination to predict the donations of the left-out participant (i.e. a leave one subject out cross-validation). We then quantified how accurately the regression predicted the donation of the left-out participants using kendall’s tau, a non-parametric estimate of correlation that is less sensitive to outliers than a parametric correlation. This multivariate approach revealed normally distributed tau values with above chance prediction accuracy (i.e. Tau>0, t_(29)_=2.365, p=0.012, BF_+0_=4.13), albeit of modest effect size (d=0.432), supporting the notion that SI does indeed contain information that relates to the magnitude of donations in the hand condition. No differences were found between the self-reported mirror synesthete and the controls (t_(28)_=0.568, p=0.575, BF_10_=0.399). Result in **Fig. S4**.

**Figure S4.**
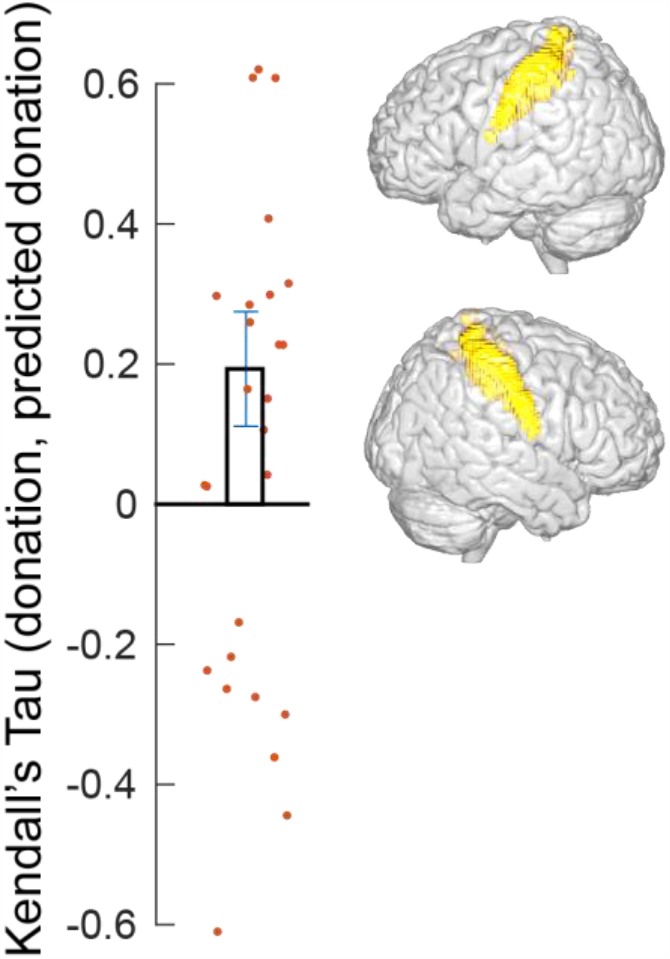
Mean (±sem) Kendall’s tau correlation between the actual donations of the participants and the ones predicted by a leave-one-out weighted partial least square regression based on parameter estimates from all SI voxels. Red dots indicate individual subject correlation. The two renders illustrate the location of the voxels included in the analysis based on the anatomy toolbox probabilistic maps of SI including BA 3a, 3b, 1 and 2.

## Notes

### Competing Interest Statement

The authors have declared no competing interest.

